# Drug-bound and -free outward-facing structures of a multidrug ABC exporter point to a swing mechanism

**DOI:** 10.1101/2021.03.12.435132

**Authors:** Vincent Chaptal, Veronica Zampieri, Benjamin Wiseman, Cédric Orelle, Juliette Martin, Kim-Anh Nguyen, Sandrine Magnard, Alexia Gobet, Margot Di Cesare, Waqas Javed, Arnaud Kilburg, Marine Peuchmaur, Julien Marcoux, Luca Monticelli, Martin Högbom, Jean-Michel Jault, Ahcène Boumendjel, Pierre Falson

## Abstract

Multidrug ABC transporters translocate drugs across membranes by a mechanism for which the molecular features of drug release are so far unknown. Here, we resolved two ATP-Mg^2+^-bound outward-facing (OF) conformations of the *Bacillus subtilis* (homodimeric) BmrA, one by X-ray crystallography without drug, and another by single-particle cryo-EM with rhodamine 6G (R6G). Two R6G molecules bind to the drug-binding cavity at the level of the outer leaflet, between transmembrane (TM) helices 1-2 of one monomer and TM5’-6’ of the other. R6G induces a rearrangement of TM1-2, highlighting a flexibility that was confirmed by H/D exchange and molecular dynamics simulations. The latter also shows a fast post-release occlusion of the cavity driven by hydrophobicity. Altogether, these data support a new swing mechanism for drug transport.

## Introduction

Multidrug ATP-Binding Cassette (ABC) exporters transport a large panel of drugs conferring a multidrug resistance (MDR) cell phenotype that leads to chemotherapy failures against pathogenic microbes and cancers. Early conceptualized (*1*), ABC exporters mainly switch between a high drug affinity inward-facing (IF) conformation in which the drug-binding pocket in the membrane domain is exposed to the inner membrane leaflet, and a low drug affinity OF conformation favoring drug release outside the cells. These proteins are made of two transmembrane domains (TMDs) typically built with twelve transmembrane helices and two nucleotide-binding domains (NBD). Drugs bind to the TMD, accessible from the inner membrane leaflet in the IF conformation. Two ATP molecules bind at the interface between the two nucleotide-binding domains (NBD) (*2, 3*), thereby stabilizing the dimer and favoring the drug occlusion that leads the reorganization of the TMD in an OF conformation (*4*).

Several exporter structures have been obtained (*5–13*), complemented with biochemical and biophysical characterizations (eg (*14–16*)), altogether contributing to a mechanistic understanding of the IF to OF transition. Moreover, the molecular mechanism by which structurally-divergent drugs bind to the IF conformation is presently better understood thanks to the structure of the human ABCB1 in complex with the anticancer drug taxol (*14*). This structure revealed that the ligand recognition is driven by the intrinsic plasticity of TM4 and TM10, required to accommodate the structure of the drug.

The question remains open as to how the structural variability of drugs is handled by those exporters to expel them and which molecular features of the protein in the OF conformation are driving this release step (*17*). So far, since the first structure released in 2006 (*5*) and almost 50 years after their discovery (*18*) no OF structure of a MDR ABC exporter with a bound drug has been solved. To that aim, the ATP-bound cryo-EM structure of ABCC1 in the presence of its substrate, Leukotriene C4, was resolved, however the location of the substrate was not determined (*19*). Previously, the crystal structure of the antibacterial peptide transporter McjD was obtained in complex with AMP-PNP and two molecules of nonyl-glucoside that were used as crystallization additive were bound in the putative drug-binding cavity (*8*). Interestingly, molecular dynamics simulation based on that structure predicted a marked flexibility of the TM1-2 region (*20*), pointing to a possible role of this region in the release of substrates. However, so far, structural information is lacking to corroborate this hypothesis, mainly due to the poor affinity of the transported substrate in the OF conformation.

Here we tackled the question by resolving two OF conformations of BmrA, a type IV ABC transporter (*21, 22*) from *B. subtilis* (*23*) conferring resistance to cervimycin C, an antibiotic produced by *Streptomyces tandæ* against Gram-positive bacteria (*24*). Using an ATPase inactive mutant, E504A (*25*), we resolved its X-ray structure in complex with ATP-Mg^2+^, which required several key steps optimization and to design specific stabilizers. We also resolved its cryo-EM structure in complex with ATP-Mg^2+^ and rhodamine 6G, a drug commonly transported by ABC exporters (*26*). Comparison of these structures enlightens how the drug binds before its release and shows how the flexibility of the TM1-2 segment drives this process, and this was confirmed by H/D exchange coupled to mass spectrometry (HDX-MS) and molecular dynamics simulations.

## Results

### Crystal structure of BmrA in OF conformation in complex with ATP-Mg^2+^

We first stabilized BmrA in its OF conformation by introducing the E504A mutation that prevents hydrolysis of ATP (*25, 27*) and substrate transport (Fig. S1) as also reported for other transporters (*14*). The protein crystallized following the procedure set-up for the mouse P-glycoprotein, using triton X-100 for extraction and a mixture of N-dodecyl-β-D-n-maltopyranoside (DDM) and cholate for purification (*28*). Quantification of detergents bound to BmrA (*29*) was helpful to produce high-quality crystals, as increasing cholate reduced the amount of DDM bound to BmrA up to 50% which proportionally reduced the estimated detergent-belt size (Fig. 1A). Diffraction patterns of the resulting crystals displayed a lattice-doubling problem that prevented their processing and which we overcame by designing a series of tailored amphiphiles **3a**-**e** (Fig. 1B; Chemistry section in Supplementary Materials) with a scaffold based on glycosyl-substituted dicarboxylates surfactants (*30*). Of note, these additives increase the thermal stability of BmrA up to ∼30 °C for **3d** (Fig. 1C, Data set 2), which helped produce better diffracting protein crystals (Fig. 1DE).

**Figure 1.**
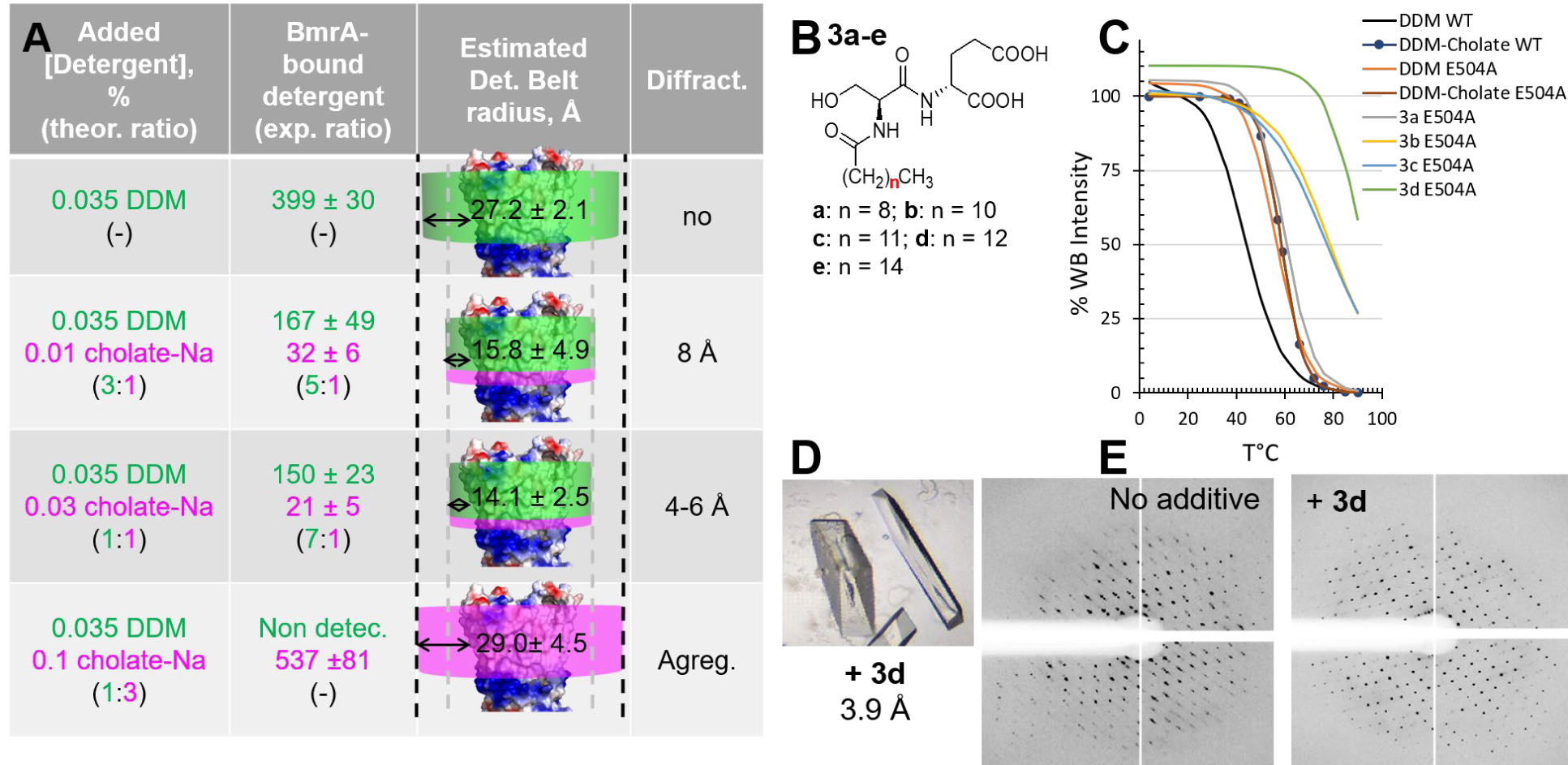
Crystallization of BmrA. **(A)** Quantification of detergents bound to BmrA. see Methods and Data set 1 for details. **(B)** Structure of the thermostabilizing amphiphilic additives. **(C)** Thermostabilisation of BmrA. For clarity, fits (2-3 independent assays) are displayed, with circles for the reference condition (DDM + cholate). Full data is provided in the Data set 2. **(D)** BmrA crystals in presence of **3d**. **(E)** Lattice problem resolution with **3d**.

We reached 3.9 Å resolution for the BmrA E504A-ATP-Mg^2+^ complex (Fig. 2, left panels; Figs. S2-S3, Table S1). Two dimers of BmrA were found in the asymmetric unit, with a r.m.s.d. of 0.7 Å over 525 residues (Fig. S2). The structure displays the characteristic type-IV fold of ABC transporters (*21*), in which the NBDs bind two ATP-Mg^2+^ in a head-to-tail mode, freezing the BmrA-E504A mutant in complex with ATP-Mg^2+^ in a typical OF conformation. The E504A mutation stabilizes the efflux pump in an OF pre-hydrolytic state, similar to the one displayed by the wild-type (WT) BmrA trapped in the transition state for ATP hydrolysis in the presence of vanadate (*31*). The extracellular side of BmrA displays an opening to a cavity likely corresponding to the drug-exit path. Importantly in the context of this study, the crystal structure shows that the loop connecting TM1 to TM2 is stabilized by few crystal contacts between half of the monomers (Fig. S2B-D), while the other half remains free of movement. All the loops nevertheless distribute around similar positions showing both the correctness of this position and the flexibility of this region (Fig. S2E).

**Figure 2.**
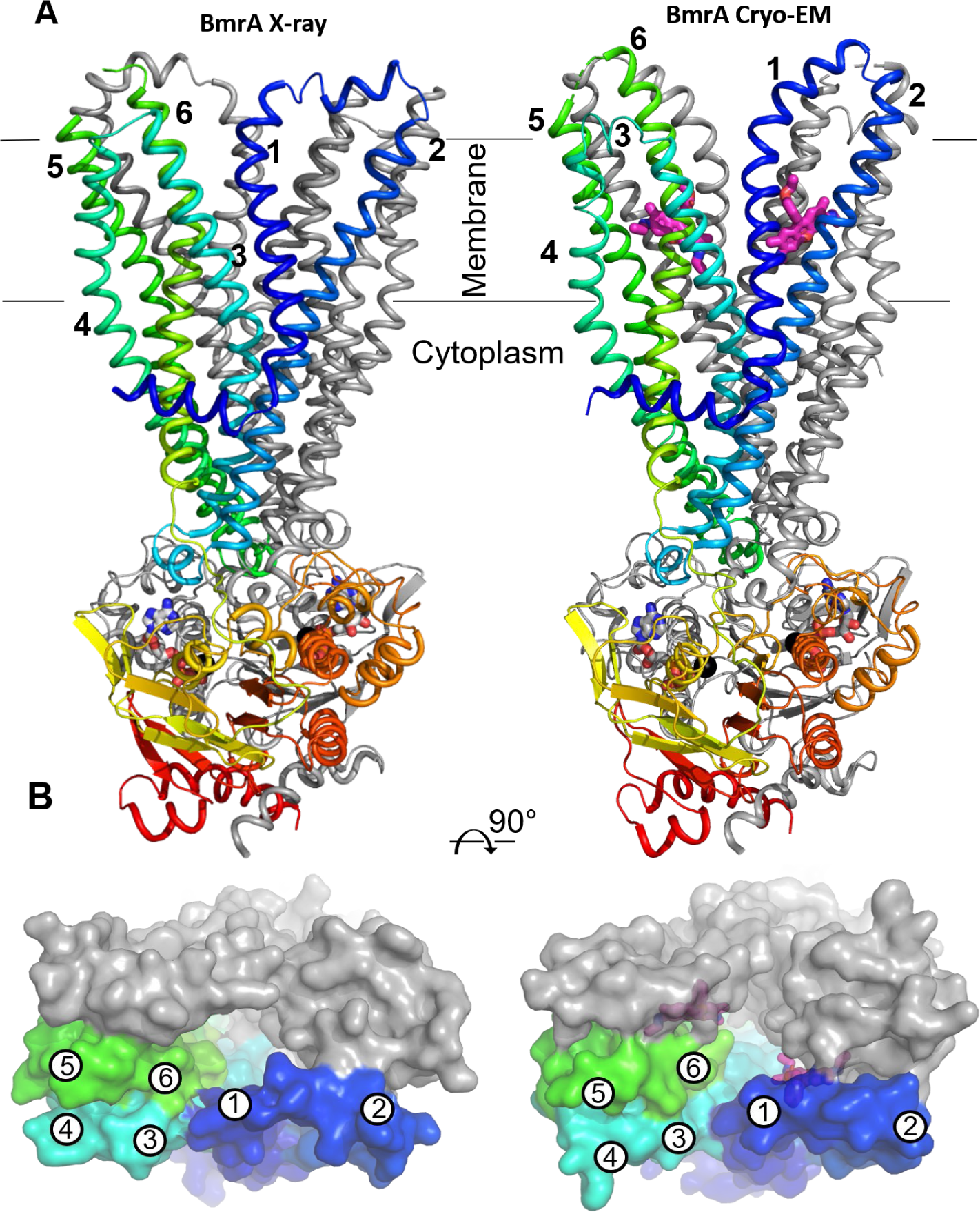
X-ray and cryo-EM structures of the BmrA E504A mutant in complex with ATP-Mg^2+^ and R6G. **(A)** Cartoons of the transporter, normal to the plane of the membrane. One monomer is in grey and the other one is rainbow colored. TM helices are numbered for the colored monomer. ATP and R6G are displayed as sticks colored by atom type and Mg^2+^ as black sphere. **(B)** Surface representation of BmrA viewed from the extracellular side to highlight the difference in the OF cavity.

### Cryo-EM structure of BmrA in the OF conformation in complex with R6G and ATP-Mg^2+^

Incubating BmrA with known ligands (*23*) gave the best crystals with R6G, which we could however not optimize beyond 5 Å (Fig. S4). This led us to move to single-particle cryo-EM (Fig. 2, Fig. S5) that allowed to build the structure using the highest resolution map using C2 symmetry. Refinement up to 3.9 Å was carried out using both sharpened and unsharpened maps, as the former lost details on the TM1-TM2 hinge movement (Fig. S6-S8). The resulting fold is very similar to that of the crystal structure with the difference in conformation of the region TM1-2 where TM1 is shifted towards TM2, resulting in a more pronounced opening of the cavity (Fig. 2B, Table S1).

We observed two additional densities in the cryo-EM density map (Fig. 3AB), seen more clearly without the application of symmetry. R6G, cholate or the polar head of DDM could be positioned in these densities, although in the case of the last two no equivalent densities was observed in the X-ray map. We therefore carried out a series of biochemical assays with the WT and E504 mutant to discriminate between these possible ligands. Both proteins were purified in DDM or DDM- cholate, followed by a reconstitution into nanodiscs (*32*), on which we probed the binding of the three compounds. We observed by intrinsic fluorescence that R6G binds to BmrA E504A purified in DDM/cholate with a 2-fold higher affinity when ATP-Mg^2+^ is present (Fig. 3C). The reverse effect was also observed, ATP-Mg^2+^ binding with 2-fold higher affinity when R6G is added (Fig. 3D). When purified in DDM, both WT and mutant BmrA displayed similar affinity for R6G (4.8 ± 0.9 and 3.3 ± 0.7 µM, *p* < 0.0001), while specific interaction could be detected neither with cholate (Fig. S9A) nor DDM (or decyl maltoside at higher concentrations), on BmrA-nanodiscs complexes (Fig. S9B). On the contrary, R6G bound to the same complexes with affinities as high as ∼0.07 ± 0.09 µM (*p* = 0.4) and ∼0.03 ± 0.02 µM (*p* = 0.1) for WT and mutant, respectively. Of note, R6G binds to nanodiscs themselves with an affinity of 3-7 µM (Fig. S9CD). These results indicate that cholate and DDM do not bind to the drug-binding site of BmrA in contrast to R6G, which binds in the sub-micromolar to micromolar range depending on the local environment. This led us to assume that this density reveals the occupancy of R6G, as displayed in Fig. 2 and Fig. 3. Finally, we evaluated the capacity of BmrA to transport R6G by quantifying the intracellular R6G level in a *B. subtilis* strain overexpressing BmrA (*24*) (BmrA+) compared to the parent strain, upon incubation with the dye (Fig. 3E). We observed that R6G accumulates ∼ 50% less in the former strain supporting that R6G is indeed exported by BmrA out of the bacteria.

**Figure 3.**
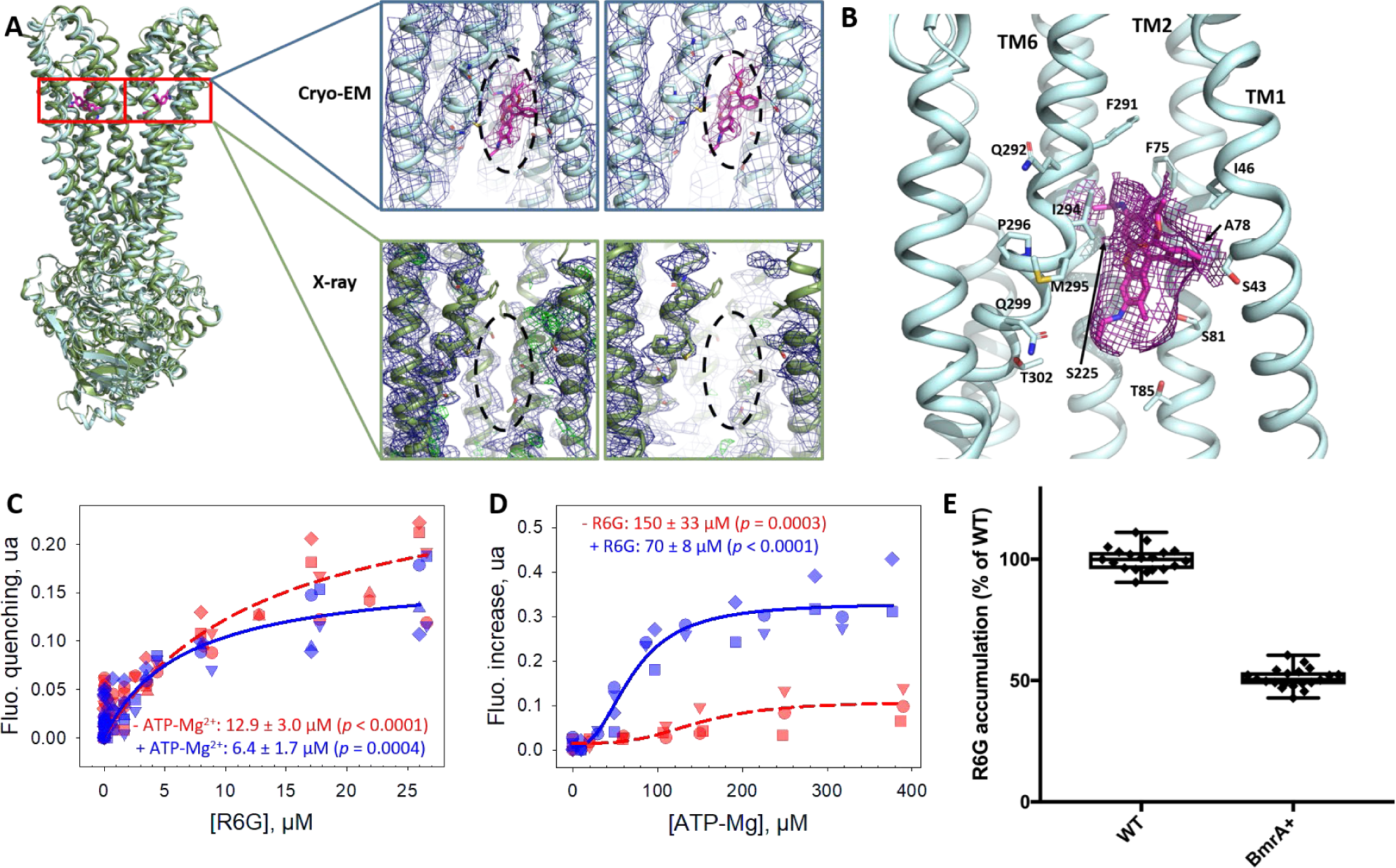
Rhodamine 6G in the cryo-EM structure of BmrA E504A mutant in complex with ATP-Mg^2+^ and its effect on BmrA activity. **(A)** Superposition of the X-ray and cryo-EM structures and zoom in the two R6G-binding sites in the cryo-EM structure compared to the X-ray structure. Density maps are shown in blue and green, respectively. R6G structure and density are in magenta. **(B)** Detail of one R6G- binding site. **(C)** Binding of R6G on BmrA E504A/DDM-cholate, with (blue) or without (red) 5 mM ATP-Mg^2+^ probed by intrinsic fluorescence. Symbols correspond to 4 independent experiments, fitted with equation 1. **(D)** Binding of ATP-Mg^2+^ with (blue) or without (red) 100 µM R6G on BmrA E504A/DDM-cholate probed by intrinsic fluorescence. Symbols correspond to 3-4 independent experiments, fitted with equation 2. **(E)** R6G accumulation in *B. subtilis* strains 168 (WT) and 8R, a mutant of *B. subtilis* 168 strain overexpressing BmrA (BmrA+), probed by fluorescence. Cells incubated with 5 µM R6G for 30 min at 37 °C were then washed, lysed, and their intracellular R6G content probed by fluorescence on supernatants, taking as reference the WT strain. Data are the average of 3 independent 3-10 replicates (*p* < 10^-10^).

The two R6G molecules bind at the level of the outward leaflet, between TM1-2 of one monomer, and TM5’-6’ of the other one. They are maintained in the cavities by a movement of TM1 towards TM2, resulting in a capped hydrophobic space sealed by residues I46, F75, L258’, M259’ and F291’. Several of these residues correspond to those found in the taxol-binding pocket of human ABCB1 (*14*) (Fig. S10).

### Structural differences between the X-ray and cryo-EM structures of BmrA highlight the mobility of the TM1-TM2 region

Although quite similar, the X-ray and Cryo-EM structures of BmrA display important local differences rendering the drug-exit path significantly different between them (Fig. 2). The differences originate from a displacement of the TM1-2 region, in the proximity of a kink starting in TM1 at residue P47, towards the end of TM1. Such displacement allows the central part of TM1 to shift from TM3 in the X-ray structure towards TM2 in the cryo-EM structure. These differences between structures solved under nearly identical conditions highlight a major local plasticity at the level of TM1-2. To evaluate its functional relevance, we compared these structures with those of previously resolved nucleotide-bound type IV ABC transporters: *E. coli*

McjD(*8, 11*), *T. thermophilus* TmrAB (*13*), human (*4*) and *C. merolæ* (*15*) ABCB1, *E. coli* MsbA (*16*), *S. aureus* Sav1866 (*5*), and *T. maritima* TM287/288 (*33*). We tentatively ranged them from the most occluded to the widest open (Fig. 4A-E). This showed that TM1 in the X-ray structure of BmrA is oriented similarly as in McjD, ABCB1 and MsbA while TM2 is shifted towards the OF conformation typically observed as in Sav1866. The loop connecting TM1 and TM2 has unwound on each side, allowing and/or accompanying the movement of TM2. In the cryo-EM structure, a consecutive displacement of TM1 shifting towards TM2 is seen, with an unwinding that takes place downward on TM1. The most open structures of Sav1866 and TM287/288 show TM1 and TM2 segments close together and separated from those forming the TM3-6 core. This motion of TM1-2 is concomitant with a wide opening of the cavity and a physical separation of the two TM3-6 cores that behave as rigid bodies. The movement would be granted by the intrinsic flexibility of ABC transporters on their external side, as hinted by the B-factors displayed in Fig. 4A. Superposing the topologically conserved regions encompassing TM3-6 core and NBD of all structures allowed to visualize the wide range of conformations of TM1-2, suggesting a hand fan motion (Fig. 4E). We could reproduce such amplitude by molecular dynamics on the present BmrA structures (Fig. 4F and detailed in the next section below). Finally, we confirmed the functional mobility of TM1-2 by probing the structural dynamics of the WT BmrA reconstituted in nanodiscs by HDX-MS experiments. We sought to identify the transmembrane regions that display an increased accessibility/flexibility when transitioning to the OF state, using BmrA either in its apo state or stabilized in its OF conformation upon vanadate-induced (Vi) nucleotide trapping (Fig. S11 and Fig. 4GH). We observed only a few transmembrane peptides that display a significantly higher deuterium uptake in the OF conformation stabilized by Vi-trapping, all localized in TM1, TM2 and TM6 (Fig. 4G). This is exemplified by the peptide 47-53 (shown by a star in panel H). Altogether, these results are consistent with the mechanical plasticity of TM1-2 inferred from the X-ray and cryo-EM structures of BmrA, which looks like a key-feature of MDR pumps allowing the release of substrates varying in size and shape.

**Figure 4.**
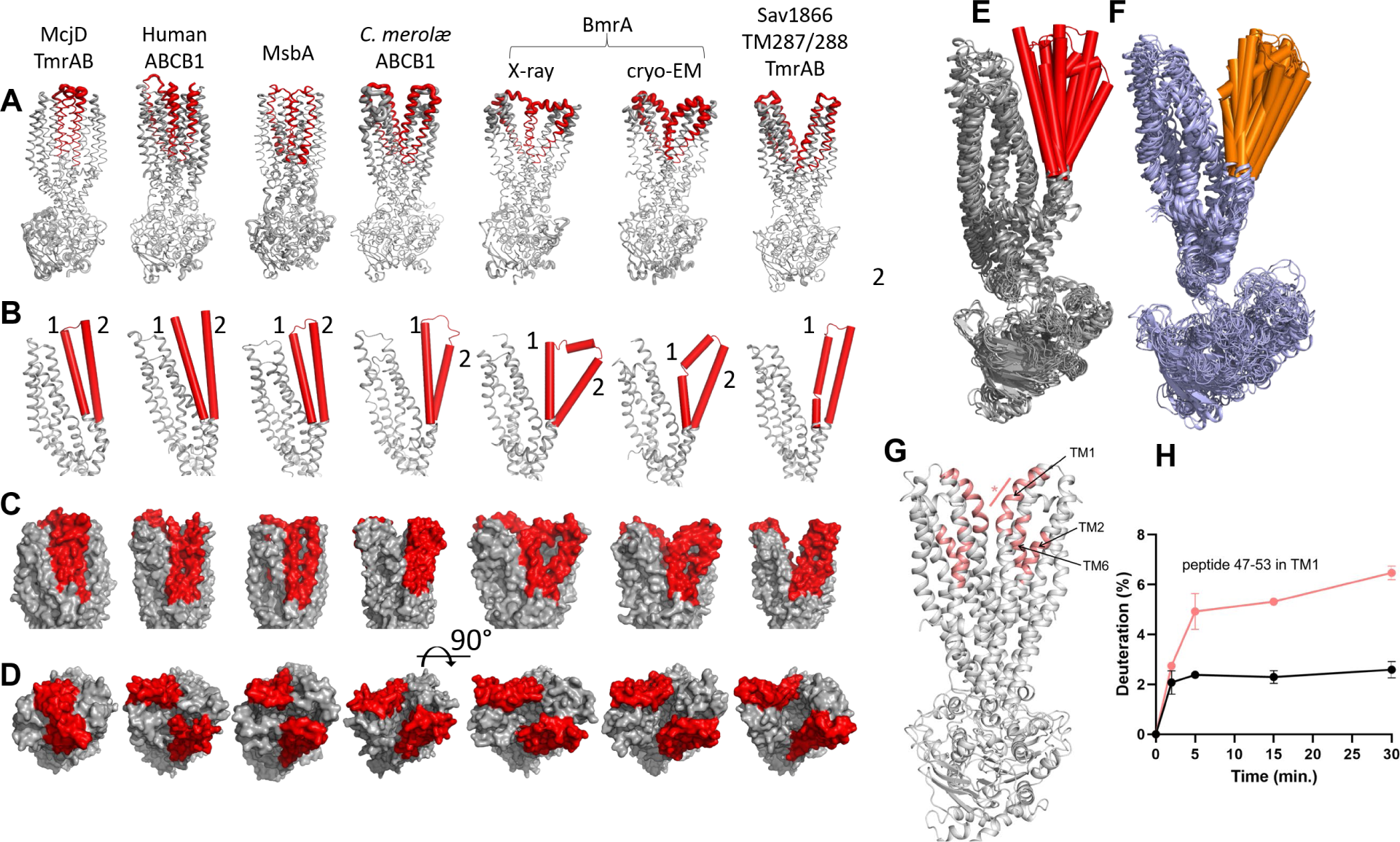
TM1-2 positions and mobility in OF BmrA and other type IV ABC transporters. **(A-D)** Views (PDB codes) of McjD (4pl0, 5ofr), TmrAB (left: 6rai, 6rak; right: 6rah-6raj), human (6c0v) and *C. merolæ* ABCB1 (6a6m), MsbA (5ttp), BmrA (this study), Sav1866 (2hyd) and TM287/288 (6qv0, 6qv1, 6qv2), superimposed from TM3 to TM6 and displayed from left to right from the most occluded to the widest open conformation. Cartoon thickness in panel **A** is proportional to B-factor. Structures are colored in grey with TM1-2 in red. **(B)** Close-up view of TM1-6 of each monomer and of the *N*-terminal half of ABCB1 with the TM1-2 segment in red cylinders. **(C)** View as in panel **B** displayed in surface. **(D)** View of the external side. **(E)** Superimposition of the different structures shown in **A**. **(F)** Molecular dynamic simulations of BmrA as detailed in Fig. 5 and corresponding section. **(G, H)** HDX-MS experiment of WT BmrA reconstituted in nanodiscs. Data were recorded after 15 min D2O exchange and only the transmembrane peptides with increased deuterium uptake in the Vi-trapped conformation as compared to the apo state are shown (salmon color, *p* < 0.01). The star indicates the position of the peptide 47-53 for which the deuterium uptake is plotted as a function of time in panel **H**, either in the Vi-trapped (salmon) or apo (black) states.

### Molecular dynamics simulations of X-ray and cryo-EM structures of BmrA

In order to get a dynamic view of the drug-exit site of BmrA, we performed all-atom molecular dynamics simulations on the present (drug-less) X-ray and cryo-EM ATP-Mg^2+^ bond structures, reconstituted in a POPE/POPG (3/1) lipid bilayer (Fig. 5A). Hence, we carried out four simulations of 500 ns on each structure in identical conditions using different starting velocities. We got an estimation of the size of the drug-exit cavity by measuring the variation with time of the perimeter formed by the C-α of residues Q52TM1 and G281_TM6_ of each monomer (Fig. 5BC). As shown, the initial perimeter of the cryo-EM structure, up to 90 Å, was larger than that of the X-ray one, up to 80 Å. Considering the simulation settings and the limited resolution of the structures we expected to observe only changes driven by strong forces. All the models undergo a closure (Fig. 5C, Fig. S12), reaching a common perimeter of ∼70 Å. BmrA shifts towards the most occluded states, as the one observed in MsbA and ABCB1 in Fig. 4. This closure is rapid, generally occurring within the initial 100 ns, as also proposed for the extracellular gate of TmrAB (*13*), and followed by large-scale structural fluctuations (Fig. 5D, Fig. S13). Of note, a closure was also obtained in simulations with longer equilibration steps (not shown). An unexpected and interesting result came from the fourth simulation generated from the X-ray structure that, by contrast to the others, rapidly opened its drug exit cavity up to ∼80 Å. This specific behavior allowed a lipid to bind between TM1-2 and 6 (left inset Fig. 5c) in a location close to that seen for rhodamine 6G, which highlights the hydrophobic nature of the ligand.

**Figure 5.**
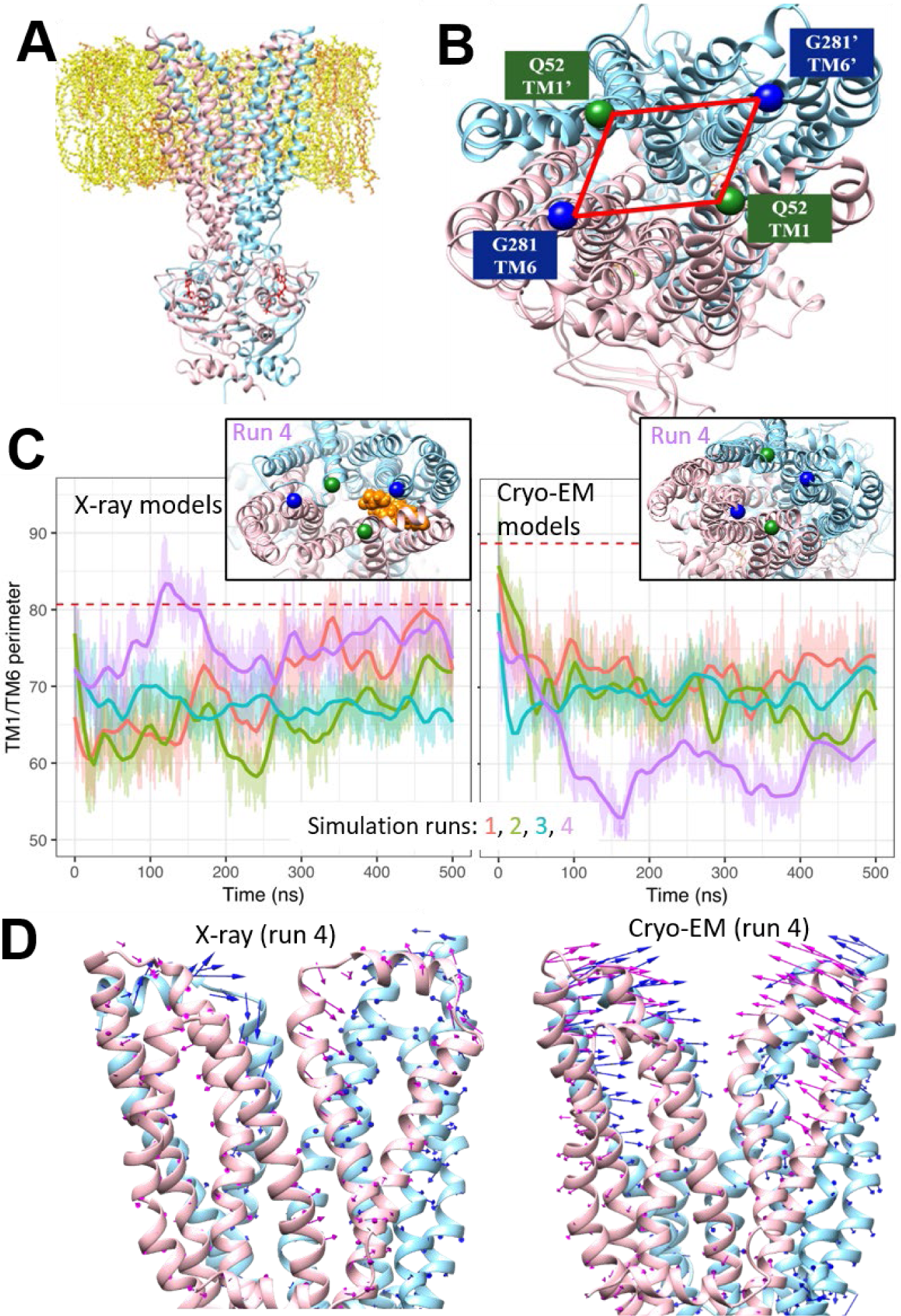
Dynamics of BmrA TMD region. **(A)** Starting model of BmrA inserted in a lipid bilayer. Chains A and B are colored in pink and cyan, ATP in red and lipids in yellow. **(B)** Close-up view showing the distance measured between residues Q52(TM1) - G281(TM6) - Q52(TM1)’ - G281(TM6)’. **(C)** Time-evolution of the (Q52-G281-Q52’-G281’) distance for each of the four simulations from X-ray and cryo-EM models. Red dashed lines indicate the initial values of the distances. Insets display snapshots of each run 4, taken at T = 336 ns and 370 ns, respectively, with on the left a lipid in orange bound to the drug-binding pocket. **(D)** Displacement of each C-α of each monomer A (pink) and B (blue) from initial X-ray (left) and cryo-EM (right) structures from the average structures.

## Discussion

The two BmrA structures presented here are the missing link in the landscape of structures of multidrug ABC transporters resolved in OF conformations and one of them reveals for the first time a structure of a type IV MDR ABC transporter with its transported substrate in a drug-release competent state. The R6G molecules are located at the level of the outer leaflet and are poised to be released from the transporter. Several parameters have contributed to stabilize thisternary complex, among them the positive effect of R6G and ATP-Mg^2+^ on their mutual affinities. The present structures reveal a drug-release site made of flexible and rigid transmembrane helices, TM1-2 and TM3-6, respectively. Combined with HDX-MS and molecular dynamics simulations, they provide new information on the mechanism of drug release from the drug-binding pocket and show how the transporter resets to an occluded conformation by closing back on itself immediately after drug release. The flexibility of multidrug ABC transporters has been well established in IF states, sampling different conformations that facilitate recognition of multiple compounds (*31, 34, 35*). The current study reveals that such a flexibility is also sampled in OF conformations, possibly with a lower amplitude. We propose that this flexibility is required to adapt the site to various substrates’ sizes, essential to secure their release to the extracellular side, and also to reset the transporter back to an occluded state, thereby preventing any trans-inhibition mechanism as reported recently (*36*). Such flexibility is also consistent with the fast dynamics of the extracellular gate of TM287/288 observed in EPR experiments (*37*) and suggests that no additional energy input is needed for drug release.

A key point arises when comparing the fast closure revealed by the simulations with the stabilized outward states observed in both X-ray and cryo-EM structures. A similar flexibility of the external part of the transmembrane segments is observed in the structures and simulations. In the case of experimental structures, the hydrophobic nature of the substrate cavity together with the accessibility to water favor its filling with amphipathic detergents as seen previously (*29*), which stabilize these OF conformations. Although detergent molecules are too mobile and flexible to be observed at these resolutions, they do however constitute excellent tools to capture such transient states. In the case of simulations in lipid membrane, the hydrophobic pocket is exposed to water, which is extremely unfavorable and leads to a rapid motion of TM1-2 that closes the pocket to shield it from water. This motion appears sufficient *per se* to reset the transporter back to an occluded conformation. The hydrophobicity of the drug-binding pocket is so important that in one simulation a single lipid molecule moved into the cavity (Fig. 5C), causing the cavity to remain open. This result fits well with the presence of a detergent molecule in the binding pocket of McjD (*8*) together with the very recent discovery of a lipid inside the structure of the Major Facilitator Superfamily protein LrmP (*38*). Altogether, this data highlights the hydrophobicity of the drug-binding pocket as the main driving force for the closing movement, independently of any ATP hydrolysis.

These findings lead us to reexamine the transport mechanism, often depicted as a cycle with a deterministic set of conformations that the transporter goes through to finally come back to its initial state. Rather, we propose a swing mechanism (Fig. 6) that relies on the flexibility of both the IF and OF conformations. The intrinsic flexibility of the exporter in the IF state allows it to sample multiple conformations. This grants the accommodation of a wide array of chemically unrelated drugs, the hallmark of multidrug transporters. Binding of ATP leads to the occluded conformation, concomitant with the plastic deformation of the outward-most part of the exporter, resulting in drug release. Here, this OF plasticity is beneficial for the release of multiple types of substrates. Hydrophobicity of the substrate binding pocket then triggers the closing of the transporter, without energy input, leading to the hypothesis that ATP hydrolysis occurs after drug release, as already proposed (*19*). The exporter thus swings back towards the IF conformation, ready for another swing. Playing together, intrinsic plasticity and hydrophobicity of the substrate binding pocket alleviate the need for precisely defined steps for transport.

**Figure 6.**
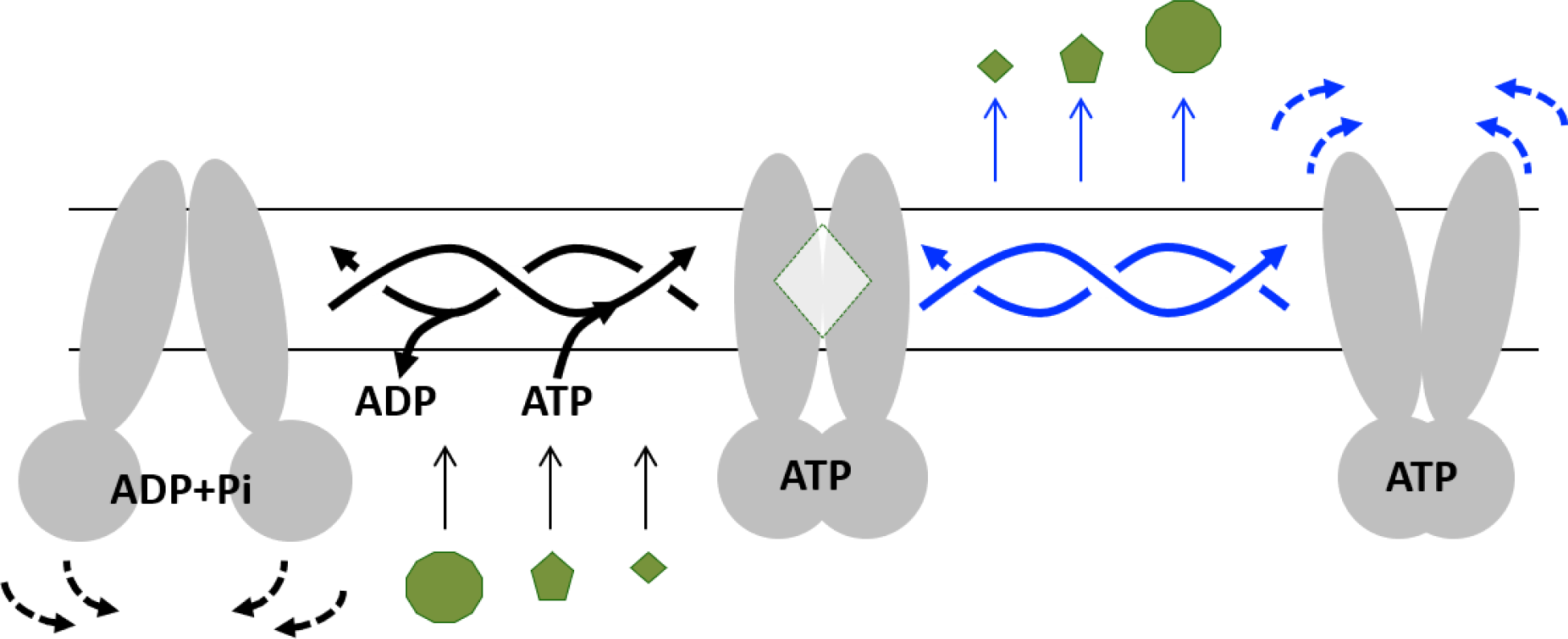
Swing mechanism of efflux. **Left**: IF conformation of the transporter displaying flexibility at the NBD level, with thus different degrees of opening of the substrate-binding cavity (black dotted arrows) and allowing for diverse shape and size of substrates. **Middle**: Occluded conformation. **Right**: OF substrate-release conformation. Release occurs *via* a plastic deformation of the external part of the transmembrane region. Deformation of the apex is adapted to the size or chemical property of the substrates (blue dotted arrows). Hydrophobicity of the drug-binding pocket triggers the immediate closing of the external part of the transporter, which swings back to the occluded state. ATP hydrolysis occurs followed by ADP and Pi release resulting in the opening in the IF conformation again, ready for another swing. Twisted arrows exemplify the different routes the transporter can take to reach any conformation, granted by the local deformability of the transmembrane helices.

## Materials and Methods

Included in the Supplementary Materials.

## Acknowledgments

**General:** We thank Pr. Klaas Martin Pos for the gift of the CD43(DE3)Δ*AcrB E. coli* strain and Pr. Hans Krügel for those of *B. subtilis* 168 and 8R. We thank Dr. Lorena Martinez for her input in the BrmA purification process in introducing the cholate-DDM mixture. We thank Synchrotron SOLEIL and ESRF staffs, the crystallogenesis platform from SFR Bioscience UMS 3444 and IBCP. We thank Drs. Clemens Von Rhein and Pierre Legrand for their help in anisotropic data processing. Cryo-EM sample screening, optimization and data collection was performed at the Cryo-EM Swedish National Facility in Stockholm, Sweden, funded by the Knut and Alice Wallenberg, Family Erling Persson and Kempe Foundations, SciLifeLab, Stockholm University, and Umeå University. BW especially thanks Marta Carroni and Julian Conrad from the Swedish National Facility in Stockholm for their technically assistance in Cryo-EM data collection

## Author contributions

VC, VZ, AK and SM purified BmrA and carried out the crystallogenesis experiments. MH granted access to cryo-EM equipment and BW carried out the cryo-EM experiments. VZ, VC and PF carried out the detergent quantifications. VC, VZ and BW resolved the structures. CO and VZ prepared the E504A mutant and carried out the transport and ATPase assays. AB and PF conceived the crystallization additives and KAN, MP, AB synthesized them. SM and PF carried out the thermostability assays. JMartin and LM carried out the dynamic simulations. WJ, MDC and JMarcoux performed the HDX-MS experiments which were supervised by J-MJ and CO. J-MJ and CO contributed to the analysis of the results. VZ, AG, MdC, SM, CO, J-MJ, VC and PF carried out the biochemical experiments. PF managed the overall project. The manuscript was written through contributions of all authors who gave their approval to its final version.

## Data and materials availability

Crystal and cryo-EM structures of BmrA-E504A have been deposited in the Protein Data Bank and Electron Microscopy Data Bank with the following codes, PDB 6r72, 6r81 and EMD-4749 (C2 symmetry), and PDB 7BG4 and EMD-12170 (no symmetry, with R6G). The HDX-MS data have been deposited to the ProteomeXchange Consortium via the PRIDE partner repository (*38*) with the data set identifier PXD022185. All data is available in the main text and the supplementary materials.

## Supplementary Materials

### Materials and Methods

#### Chemistry

Solvents and reagents were purchased from commercial sources and used without further purification. Reactions were monitored by thin layer chromatography (TLC) using commercial silica gel 60 F_254_ coated plates from Macherey-Nagel. Visualization were carried out under UV light at 254 and 365 nm and/or heating with a solution of sulfuric acid/acetic acid/water or phosphomolybdic acid/cerium sulfate/sulfuric acid/water or ninhydrin stain or iodine vapor. Purifications were performed by gravity column chromatography using silica gel 60 (230-400 mesh) from Macherey-Nagel or by automatic Reveleris® X2 flash chromatography system. MPs were measured using a Büchi B540 melting point apparatus and are uncorrected. Electrospray ionization (ESI) mass spectra were obtained on an Esquire 3000 Plus Bruker Daltonis instrument with a nanospray inlet. Accurate mass measurements (HRMS) were carried out on an ESI/QTOF with the Waters Xevo G2-S QTof device. Analyses were performed by the analytical service of *Institut de Chimie Moléculaire de Grenoble* (ICMG). Spectra were recorded in deuterated solvents on Bruker Avance spectrometers at 400 or 500 MHz for ^1^H and 100 or 125 MHz for ^13^C NMR, respectively. Chemical shifts (*δ*) are reported in parts per million (ppm) relative to the solvent [^1^H: δ(acetone-*d_6_*) = 2.05 ppm, δ(DMSO-*d_6_*) = 2.50 ppm, δ(CD_3_OD) = 3.31 ppm, δ(CDCl_3_) = 7.26 ppm; ^13^C: δ(DMSO-*d_6_*) = 39.5 ppm, δ(CD_3_OD) = 49.0 ppm, δ(CDCl_3_) = 77.2 ppm, δ(acetone-*d_6_*) = 206.3 ppm]. Multiplicity of signals is reported as followed: s (singlet), bs (broad singlet), d (doublet), t (triplet), q (quartet), qt (quintet), st (septet), dd (doublet of doublet), ddd (doublet of doublet of doublet), dt (doublet of triplet) ddt (doublet of doublet of triplet) and m (multiplet). Coupling constants (J) are given in Hertz (Hz). When direct signal assignments were difficult, additional spectra were acquired (J-mod, COSY, HMQC or HMBC).

#### Synthesis of amphiphiles 3a-3e as crystallization additives

Crystallization additives were obtained according to the synthetic scheme shown below.

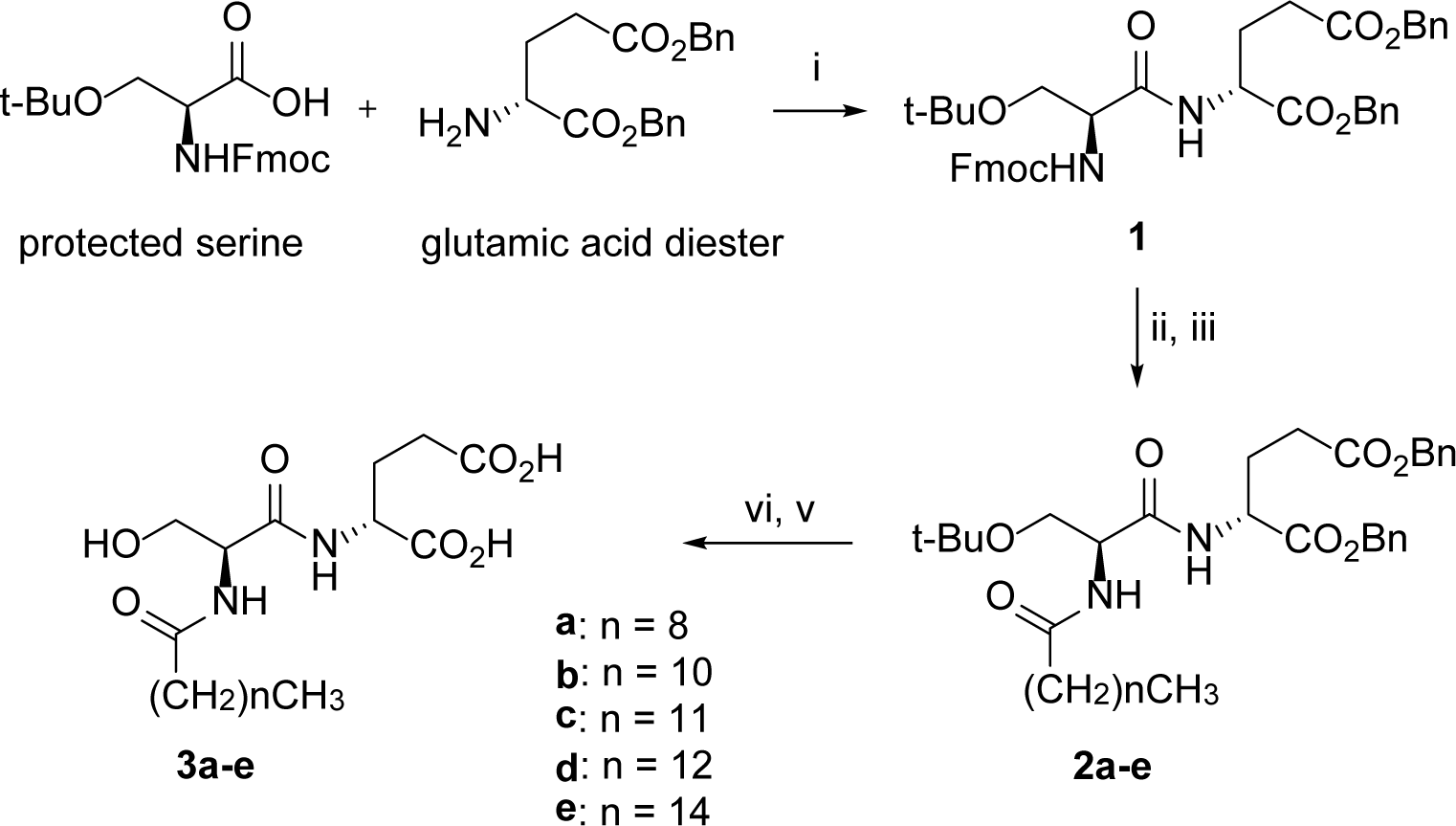

Reagents and Conditions. **i**. TBTU, DIEA, DMF; **ii**. Et_2_NH, CH_2_Cl_2_; **iii**. R-CO-Cl, DMAP, pyridine, CH_2_Cl_2_; **vi**. H_2_, Pd/C, MeOH; **v**. TFA, CH_2_Cl_2_. Synthesis of compound 1. Dibenzyl (R)-2-{(S)-2-[((9H-fluoren-9-yl)methoxycarbonyl) amino]-3- tert-butoxypropanamido} glutarate.

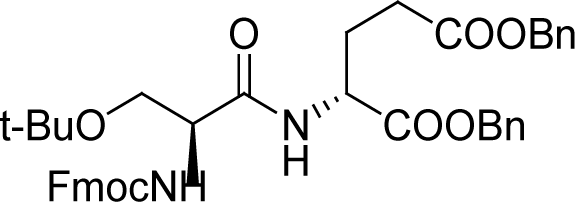

To a solution of protected serine (3.5 g, 9.12 mmol) in anhydrous DMF (15 mL/mmol) were successively added the glutamic acid diester (9.0 g, 18.24 mmol, 2 equiv.), TBTU (1.2 equiv.) and DIPEA (5 equiv.). The mixture was stirred at room temperature (rt) under N_2_ atmosphere for 3 h. After completion of the reaction, water (15 mL/mmol) was added. The compound precipitated and was crystallized in a mixture of CH_2_Cl_2_/Et_2_O to provide compound 1 (5.14 g, 81% yield). R*f* = 0.50 (cyclohexane/EtOAc 7:3); MP = 126-128 ^°^C; ^1^H NMR (400 MHz, CDCl_3_) δ ppm 1.19 (s, 9H), 1.99-2.11 (m, 1H), 2.22-2.35 (m, 1H), 2.32-2.55 (m, 2H), 3.41 (dd, *J* = 8.3, 8.3 Hz, 1H), 3.75-3.87 (m, 1H), 4.23 (t, *J* = 7.1 Hz, 1H), 4.24-4.33 (m, 1H), 4.40 (d, *J* = 6.8 Hz, 2H), 4.68-4.76 (m, 1H), 5.09 (s, 2H), 5.17 (s, 2H), 5.78 (bs, 1H), 7.21-7.46 (m, 15H), 7.61 (d, *J* = 7.0 Hz, 2H), 7.76 (d, *J* = 7.5 Hz, 2H). _13_C NMR (100 MHz, CDCl_3_) δ ppm 27.4 (3xCH_3_), 27.5 (CH_2_), 30.0 (CH_2_), 47.1 (CH), 51.8 (CH), 54.6 (CH), 61.7 (CH_2_), 66.5 (CH_2_), 67.2 (CH_2_), 67.4 (CH_2_), 74.3

(C), 120.0 (2xCH), 125.1 (2xCH), 127.1 (2xCH), 127.7 (2xCH), 128.2-128.7 (10xCH), 135.1 (C), 135.7 (C), 141.3 (2xC), 143.7 (2xC), 156.1 (C), 170.1 (C), 171.2 (C), 172.3 (C); MS (ESI+) m/z (%) 426 (100), 570 (3), 715 (1) [M+Na]_+_; HRMS (ESI+) m/z, calculated for C_41_H_45_N_2_O_8_ 693.3176, found 693.3156.

#### Synthesis of compounds 2

*Fmoc deprotection.* To a solution of compound **1** (1 equiv.) in anhydrous dichloromethane (20 mL/mmol) was added diethylamine (20 equiv.). The reaction mixture was stirred at rt under N_2_ atmosphere overnight. The volatiles were removed under reduced pressure. To eliminate the residual diethylamine, the crude product was diluted in dichloromethane, washed with a saturated sodium bicarbonate (NaHCO_3_) solution, dried over MgSO_4_, filtered and concentrated under reduced pressure and used for the next steps without further purification.

*Amide formation.* The crude compound obtained in the previous step (1 equiv.) was dissolved in anhydrous dichloromethane (30 mL/mmol). The acyl chloride derivative was added (2 equiv.), together with dimethylaminopyridine (DMAP) (0.5 equiv.) and pyridine (34 equiv.). The reaction mixture was stirred at rt under N_2_ atmosphere overnight. The reaction mixture was acidified to pH = 3 with an aqueous solution of HCl 10% and extracted with dichloromethane. The combined organic layers were washed with brine and dried over MgSO_4_, filtered, and concentrated under reduced pressure. The crude product was purified by silica gel column chromatography. **Compound 2a**. Dibenzyl (R)-2-[(S)-3-tert-butoxy-2-(decanamido)propanamido]glutarate

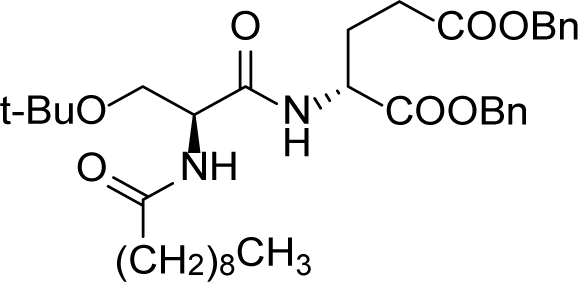

The crude product was prepared starting from **1** (500 mg, 0.70 mmol) and commercially available decanoyl chloride (267 mg, 1.40 mmol). After purification by column chromatography on silica gel (cyclohexane/EtOAc 8:2 to 7:3), the pure product **2a** (195 mg, 0.31 mmol, 45%) was obtained as a white solid. R*f* = 0.18 (cyclohexane/EtOAc 7:3). MP = 82-85 °C; _1_H NMR (400 MHz, CDCl_3_) *δ* ppm 0.87 (t, *J* = 6.9 Hz, 3H), 1.16 (s, 9H), 1.19-1.35 (m, 12H), 1.55-1.66 (m, 2H), 1.96-2.07 (m, 1H), 2.21 (t, *J* = 7.8 Hz, 2H), 2.21-2.30 (m, 1H), 2.32-2.50 (m, 2H), 3.31 (dd, *J* = 8.7, 8.7 Hz, 1H), 3.80 (dd, *J* = 8.7, 4.2 Hz, 1H), 4.44-4.50 (m, 1H), 4.65-4.72 (m, 1H), 5.08 (s, 2H), 5.15 (s, 2H), 6.40 (d, *J* = 6.4 Hz, 1H, NH), 7.27-7.36 (m, 11H) ; _13_C NMR (100 MHz, CDCl_3_) *δ* ppm 14.1 (CH_3_), 22.7 (2xCH_2_), 25.6 (CH_2_), 27.4 (3xCH_3_), 27.5 (CH_2_), 29.3-29.44 (4xCH_2_), 30.0 (CH_2_), 31.9 (CH_2_), 36.6 (CH_2_), 51.8 (CH), 53.0 (CH), 61.3 (CH_2_), 66.5 (CH_2_), 67.3 (CH_2_), 74.3 (C), 128.3-128.7 (10xCH), 135.2 (C), 135.8 (C), 170.4 (C), 171.2 (C), 172.3 (C), 173.3 (C) ; MS (ESI+) *m/z* (%) 626 (30) [M+H]_+_, 648 (100) [M+Na]_+_; HRMS (ESI+) *m/z*, calculated for C_36_H_53_N_2_O_7_ 625.3853, found 625.3846.

**Compound 2b**. Dibenzyl (R)-2-[(S)-3-tert-butoxy-2-(dodecanamido)propanamido]glutarate

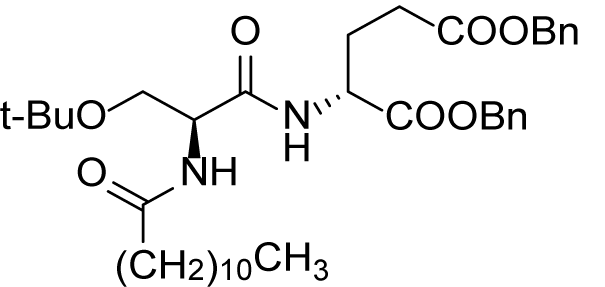

The crude product was prepared starting from **1** (400mg, 0.58 mmol) and commercially available dodecanoyl chloride (252 mg, 1.15 mmol). After purification by column chromatography on silica gel (cyclohexane/EtOAc 8:2 to 7:3), the pure product **2b** (182 mg, 0.28 mmol, 48%) was obtained as a white solid. R*f* = 0.12 (cyclohexane/EtOAc 8:2) ; MP **=** 67-69 °C ; _1_H NMR (400 MHz, CDCl_3_) *δ* ppm 0.88 (t, *J* = 6.9 Hz, 3H), 1.17 (s, 9H), 1.19-1.35 (m, 16H), 1.56-1.67 (m, 2H), 1.96-2.08 (m, 1H), 2.21 (t, *J =* 7.6 Hz, 2H), 2.21-2.32 (m, 1H), 2.32-2.50 (m, 2H), 3.30 (dd, *J =* 8.7, 8.7 Hz, 1H), 3,81 (dd, *J* =

8.7, 4.2 Hz, 1H), 4.42-4.49 (m, 1H), 4.64-4.72 (m, 1H), 5.09 (s, 2H), 5.16 (s, 2H), 6.38 (d, *J =* 6.3 Hz, 1H, NH), 7.23-7.39 (m, 11H); _13_C NMR (100 MHz, CDCl3) *δ* ppm 14.2 (CH3), 22.8 (CH_2_),

25.6 (CH_2_), 27.5 (3xCH_3_), 27.6 (CH_2_), 29.4-29.7 (6xCH_2_), 30.0 (CH_2_), 32.0 (CH_2_), 36.7 (CH_2_), 51.9 (CH), 53.1 (CH), 61.4 (CH_2_), 66.6 (CH_2_), 67.5 (CH_2_), 74.4 (C), 128.4-128.8 (10xCH), 135.2 (C), 135.9 (C), 170.4 (C), 171.3 (C), 172.4 (C), 173.4 (C) ; MS (ESI+) *m/z* (%) 131 (30), 199 (40), 654 (50) [M+H]^+^, 677 (100), 699 (20); HRMS (ESI+) *m/z*, calculated for C_38_H_57_N_2_O_7_ 653.4166, found 653.4158.

**Compound 2c**. Dibenzyl (R)-2-[(S)-3-tert-butoxy-2-(tridecanamido)propanamido]glutarate.

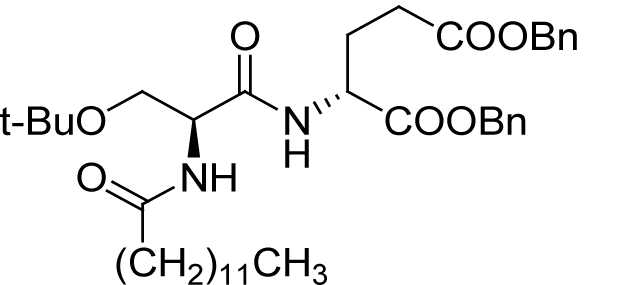

The crude product was prepared starting from **1** (500 mg, 0.70 mmol) and commercially available tridecanoyl chloride (326 mg, 1.40 mmol). After purification by column chromatography on silica gel (cyclohexane/EtOAc 8:2 to 7:3), the pure product **2c** (233 mg, 0.35 mmol, 50%) was obtained as a white solid.

R*f* = 0.24 (cyclohexane/EtOAc 7:3); MP **=** 68-71 °C ; _1_H NMR (400 MHz, CDCl_3_) *δ* ppm 0.88 (t, *J* = 6.9 Hz, 3H), 1.17 (s, 9H), 1.20-1.36 (m, 18H), 1.55-1.67 (m, 2H), 1.96-2.08 (m, 1H), 2.21 (t, *J =* 7.6 Hz, 2H), 2.21-2.32 (m, 1H), 2.32-2.50 (m, 2H), 3.31 (dd, *J =* 8.7, 8.7 Hz, 1H), 3.80 (dd, *J* = 8.7, 4.2 Hz, 1H), 4.44-4.50 (m, 1H), 4.63-4.73 (m, 1H), 5.08 (s, 2H), 5.15 (s, 2H), 6.41 (d, *J =* 6.4 Hz, 1H, NH), 7.25-7.37 (m, 11H); _13_C NMR (100 MHz, CDCl_3_) *δ* ppm 14.1 (CH_3_), 22.7 (CH_2_), 25.5 (CH_2_), 27.4 (3xCH_3_), 27.5 (CH_2_), 29.3-29.7 (7xCH_2_), 29.9 (CH_2_), 31.9 (CH_2_), 36.5 (CH_2_), 51.8 (CH), 53.0 (CH), 61.3 (CH_2_), 66.5 (CH_2_), 67.3 (CH_2_), 74.2 (C), 128.3-128.6 (10xCH), 135.2 (C), 135.8 (C), 170.4 (C), 171.2 (C), 172.3 (C), 173.3 (C) ; MS (ESI+) *m/z* (%) 668 (20) [M+H]^+^, 690 (100) [M+Na]^+^; HRMS (ESI+) *m/z*, calculated for C_39_H_59_N_2_O_7_ 667.4322, found 667.4334.

**Compound 2d**. Dibenzyl (R)-2-[(S)-3-t-butoxy-2-(tetradecanamido)propanamido] glutarate.

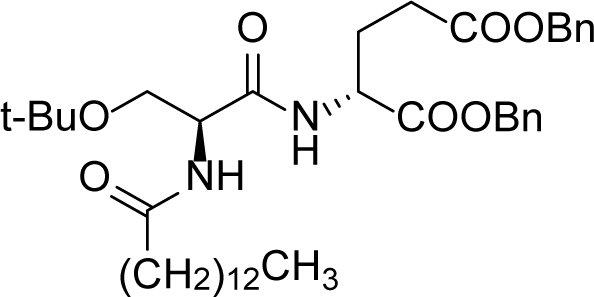

The crude product was prepared starting from **1** (400 mg, 0.58 mmol) and commercially available tetradecanoyl chloride (285 mg, 1.15 mmol). After purification by column chromatography on silica gel (cyclohexane/EtOAc 8:2 to 7:3), the pure product **2d** (187 mg, 0.27 mmol, 48%) was obtained as a white solid. R*f* = 0.07 (cyclohexane/EtOAc 8:2); MP = 71-73 °C ; ^1^H NMR (400 MHz, CDCl_3_) *δ* ppm 0.88 (t,

*J* = 6.9 Hz, 3H), 1.17 (s, 9H), 1.20-1.36 (m, 20H), 1.56-1.67 (m, 2H), 1.97-2.08 (m, 1H), 2.21 (t, *J* = 7.6 Hz, 2H), 2.23-2.50 (m, 3H), 3.29 (dd, *J =* 8.7, 8.7 Hz, 1H), 3.80 (dd, *J* = 8.7, 4.2 Hz, 1H),

4.41-4.49 (m, 1H), 4.63-4.72 (m, 1H), 5.09 (s, 2H), 5.14 (d, *J =* 12.3 Hz, 1H), 5.18 (d, *J =* 12.3 Hz, 1H), 7.25-7.39 (m, 11H); ^13^C NMR (100 MHz, CDCl_3_) *δ* ppm 14.2 (CH_3_), 22.8 (CH_2_), 25.6 (CH_2_), 27.5 (3xCH_3_), 27.6 (CH_2_), 29.4-29.8 (8xCH_2_), 30.0 (CH_2_), 32.0 (CH_2_), 36.7 (CH_2_), 51.9 (CH), 53.1 (CH), 61.4 (CH_2_), 66.6 (CH_2_), 67.5 (CH_2_), 74.4 (C), 128.4-128.8 (10xCH), 135.2 (C), 135.8 (C), 170.5 (C), 171.3 (C), 172.4 (C), 173.4 (C) ; MS (ESI+) *m/z* (%) 131 (65), 199 (100), 682 (60) [M+H]^+^ ; HRMS (ESI+) *m/z* calculated for C_40_H_61_N_2_O7 681,4479, found 681,4447.

**Compound 2e**. Dibenzyl (R)-2-[(S)-3-tert-butoxy-2-(hexadecanamido)propanamido]glutarate

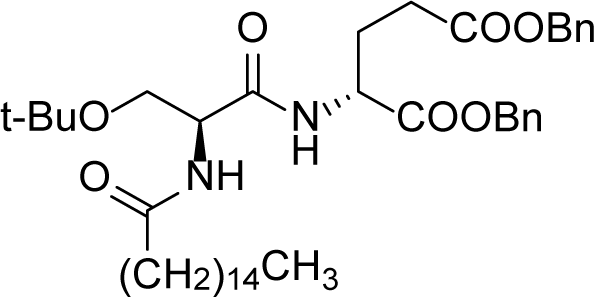

The crude product was prepared starting from **1** (300 mg, 0.43 mmol) and commercially available hexadecanoyl chloride (238 mg, 0.87 mmol). After purification by column chromatography on silica gel (cyclohexane/EtOAc 8:2), the pure product **2e** (93 mg, 0.13 mmol, 30%) was obtained as a white solid.

R*f* = 0.11 (8:2 cyclohexane/EtOAc). MP = 69-71 °C. ^1^H NMR (400 MHz, CDCl_3_) *δ* ppm 0.88 (t, *J* = 6.8 Hz, 3H), 1.17 (s, 9H), 1.20-1.37 (m, 24H), 1.55-1.68 (m, 2H), 1.97-2.09 (m, 1H), 2.21 (t, *J* = 7.6 Hz, 2H), 2.16-2.51 (m, 3H), 3.29 (dd, *J =* 8.7, 8.7 Hz, 1H), 3.81 (dd, *J* = 8.7, 4.2 Hz, 1H), 4.39-4.49 (m, 1H), 4.62-4.72 (m, 1H), 5.09 (s, 2H), 5.14 (d, *J =* 12.3 Hz, 1H), 5.18 (d, *J =* 12.3 Hz, 1H), 6.36 (d, *J =* 6.3 Hz, 1H), 7.20-7.40 (m, 11H); _13_C NMR (100 MHz, CDCl_3_) *δ* ppm 14.3 (CH_3_), 22.8 (CH_2_), 25.7 (CH_2_), 27.5 (3xCH_3_), 27.6 (CH_2_), 29.4-29.8 (10xCH_2_), 30.1 (CH_2_), 32.1 (CH_2_), 36.7 (CH_2_), 51.9 (CH), 53.1 (CH), 61.4 (CH_2_), 66.6 (CH2), 67.5 (CH2), 74.5 (C), 128.4- 128.8 (10xCH), 135.2 (C), 135.9 (C), 170.5 (C), 171.3 (C), 172.4 (C), 173.5 (C); MS (ESI+) *m/z* (%) 199 (15), 710 (100) [M+H]^+^, 732 (15) [M+Na]^+^; HRMS (ESI+) *m/z* calculated for C_42_H_65_N_2_O_7_ 709.4792 [M+H]^+^, found 709.4805.

#### Synthesis of compounds 3

*Catalytic hydrogenolysis.* To a degassed solution of a compound **2** (1 equiv.) in MeOH (100 mL/mmol) was added Pd/C 10% (200 mg/mmol). The reaction mixture was stirred at rt under H_2_ atmosphere from 4 h to overnight. After filtration over Celite® to remove the catalyst, the solvent was evaporated under reduced pressure. The residue was used directly for the next step or washed with cyclohexane and/or dichloromethane to obtain the product which was used as is for the *t*-Bu deprotection step.

*t-Butyl deprotection.* To a solution of *t*-Bu-intermediate, obtained in the previous step (1 equiv.) in anhydrous dichloromethane (12 mL/mmol) at 0 °C was added dropwise TFA (4 mL/mmol). The reaction mixture was stirred at rt under N_2_ atmosphere overnight. The volatiles were removed under reduced pressure and the residue was dissolved in DCM. A solution of NaOH (2 M) was added to pH = 11-12. The aqueous layer was washed with EtOAc before being acidified to pH 1-2 with concentrated HCl and extracted 3 times with EtOAc. The combined organic layers were dried over MgSO_4_, filtered and concentrated under reduced pressure. The residue was washed with DCM to obtain the pure product.

**Compound 3a**. (R)-2-[(S)-2-(Decanamido)-3-hydroxypropanamido]glutaric acid.

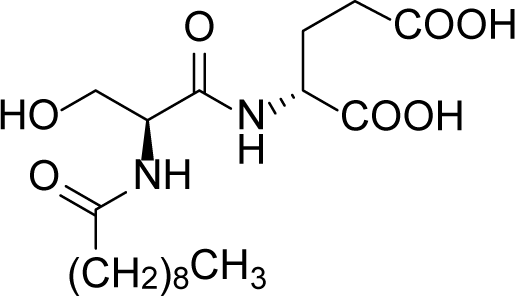

The pure product (white solid, 182 mg, 0.47 mmol, 94%) was prepared starting from **2a** (310 mg, 0.50 mmol). MP = 53-57 °C; _1_H NMR (400 MHz, CD_3_OD) *δ* ppm 0.89 (t, *J =* 6.8 Hz, 3H), 1.22-1.38 (m, 12H), 1.56-1.69 (m, 2H), 1.90-2.03 (m, 1H), 2.14-2.26 (m, 1H), 2.29 (t, *J =* 7.3 Hz, 2H),2.36-2.43 (m, 2H), 3.73-3.84 (m, 2H), 4.42-4.52 (m, 2H); _13_C NMR (100 MHz, CD_3_OD) *δ* ppm14.4 (CH_3_), 23.7 (CH_2_), 26.8 (CH_2_), 27.9 (CH_2_), 30.3-30.5 (4xCH_2_), 31.1 (CH_2_), 33.0 (CH_2_), 36.9 (CH_2_), 53.3 (CH), 56.6 (CH), 63.1 (CH_2_), 172.5 (C), 174.9 (C), 176.5 (C), 176.5 (C); MS (ESI-) *m/z* (%) 387 (100) [M-H]_-_, 404 (20); HRMS (ESI-) *m/z* calculated for C_18_H_31_N_2_O_7_ 387.2131 [M- H]^-^, found 387.2140.

**Compound 3b.** (R)-2-[(S)-2-(Dodecanamido)-3-hydroxypropanamido]glutaric acid.

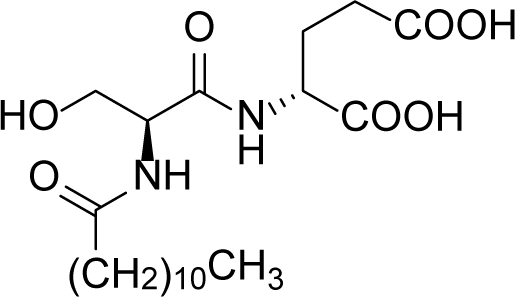

The pure product (white solid, 1.68 g, 4.04 mmol, 70%) was prepared starting from **2b** (3.76 g, 5.80 mmol). MP = 100-102 °C; ^1^H NMR (400 MHz, CD_3_OD) *δ* ppm 0.90 (t, *J =* 6.9 Hz, 3H), 1.22-1.38 (m, 16H), 1.57-1.67 (m, 2H), 1.91-2.01 (m, 1H), 2.15-2.26 (m, 1H), 2.29 (t, *J =* 7.2 Hz, 2H), 2.37-2.44 (m, 2H), 3.72-3.81 (m, 2H), 4.43-4.51 (m, 2H); _13_C NMR (100 MHz, CD_3_OD) *δ* ppm 14.4 (CH_3_), 23.7 (CH_2_), 26.8 (CH_2_), 27.9 (CH_2_), 30.4-30.7 (6xCH_2_), 31.0 (CH_2_), 33.1 (CH_2_), 36.9 (CH_2_), 53.1 (CH), 56.6 (CH), 63.1 (CH_2_), 172.6 (C), 174.6 (C), 176.4 (C), 176.5 (C); MS (ESI-) *m/z* (%) 157 (40), 199 (30), 387 (80), 415 (100) [M-H]^-^; HRMS (ESI-) *m/z* calculated for C_20_H_35_N_2_O_7_ 415.2444 [M-H]^-^, found 415.2447.

**Compound 3c**. (R)-2-[(S)-2-(Tridecanamido)-3-hydroxypropanamido]glutaric acid.

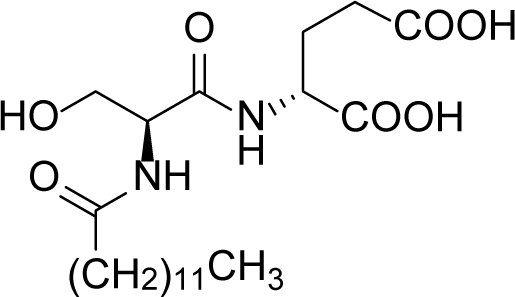

The pure product (white solid, 115 mg, 0.27 mmol, 89%) was prepared starting from **2c** (198 mg, 0.30 mmol). MP = 58-63 °C; ^1^H NMR (400 MHz, CD_3_OD) *δ* ppm 0.89 (t, *J* = 6.9 Hz, 3H), 1.21- 1.41 (m, 18H), 1.54- 1.68 (m, 2H), 1.90-2.04 (m, 1H), 2.14-2.27 (m, 1H), 2.29 (t, *J =* 7.4 Hz, 2H), 2.36-2.48 (m, 2H), 3.71-3.83 (m, 2H), 4.43-4.53 (m, 2H); _13_C NMR (100 MHz, CD_3_OD) *δ* ppm 14.4 (CH_3_), 23.7 (CH_2_), 26.8 (CH_2_), 27.8 (CH_2_), 30.3-30.9 (7xCH_2_), 31.0 (CH_2_), 33.0 (CH_2_), 36.9 (CH_2_), 53.1 (CH), 56.6 (CH), 63.1 (CH2), 172.6 (C), 174.6 (C), 176.4 (C), 176.5 (C); MS (ESI-) *m/z* (%) 429 (100) [M-H]^-^, 446 (30); HRMS (ESI-) *m/z* calculated for C_21_H_37_N_2_O_7_ 429.2601 [M-H]^-^, found 429.2599.

**Compound 3d**. (R)-2-[(S)-2-(Tetradecanamido)-3-hydroxypropanamido]glutaric acid

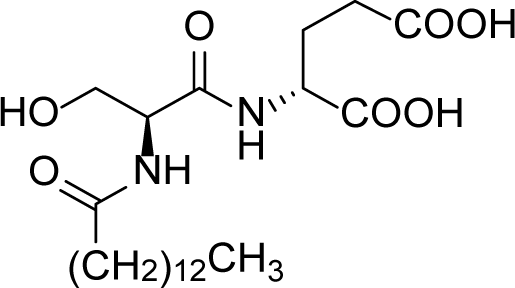

The pure product (white solid, 57 mg, 0.13 mmol, quantitative) was prepared starting from **2d** (87 mg, 0.13 mmol). MP = 109-112 °C; ^1^H NMR (400 MHz, CD_3_OD) *δ* ppm 0.90 (t, *J* = 6.8 Hz, 3H),1.19-1.39 (m, 20H), 1.57- 1.67 (m, 2H), 1.90-2.02 (m, 1H), 2.16-2.26 (m, 1H), 2.29 (t, *J =* 7.4 Hz, 2H), 2.36-2.45 (m, 2H), 3.71-3.84 (m, 2H), 4.43-4.52 (m, 2H); _13_C NMR (100 MHz, CD_3_OD) *δ* ppm 14.4 (CH_3_), 23.7 (CH_2_), 26.8 (CH_2_), 27.9 (CH_2_), 30.4-30.9 (8xCH_2_), 31.1 (CH_2_), 33.0 (CH_2_), 36.9 (CH_2_), 53.3 (CH), 56.6 (CH), 63.1 (CH_2_), 172.6 (C), 174.9 (C), 176.5 (C), 176.5 (C); MS (ESI-) *m/z* (%) 443 (100) [M-H]_-_; HRMS (ESI-) *m/z* calculated for C_22_H_39_N_2_O_7_ 443.2757 [M- H]^-^, found 443.2754.

**Compound 3e.** (R)-2-[(S)-2-(Hexadecanamido)-3-hydroxypropanamido]glutaric acid

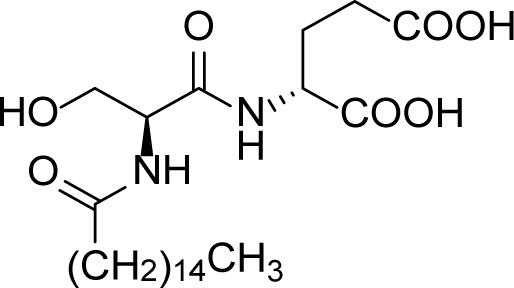

The pure product (white solid, 36 mg, 0.08 mmol, 65%) was prepared starting from **2e** (83 mg, 0.18 mmol). MP = 111-113 °C; ^1^H NMR (400 MHz, CD_3_OD) *δ* ppm 0.90 (t, *J* = 6.8 Hz, 3H), 1.19-1.37 (m, 24H), 1.55-1.68 (m, 2H), 1.90-2.03 (m, 1H), 2.15-2.26 (m, 1H), 2.29 (t, *J =* 7.4 Hz, 2H), 2.36-2.44 (m, 2H), 3.72-3.82 (m, 2H), 4.43-4.52 (m, 2H); _13_C NMR (100 MHz, CD_3_OD) *δ* ppm 14.4 (CH_3_), 23.7 (CH_2_), 26.8 (CH_2_), 27.9 (CH_2_), 30.4-30.8 (10xCH_2_), 31.1 (CH_2_), 33.1 (CH_2_), 36.9 (CH_2_), 53.2 (CH), 56.6 (CH), 63.1 (CH_2_), 172.6 (C), 174.7 (C), 176.5 (C), 176.5 (C); MS (ESI) *m/z* (%) 471 (100) [M-H]_-_; HRMS (ESI-) *m/z* calculated for C_24_H_43_N_2_O_7_ 471.3070, found 471.3057.

#### Biochemistry

##### Products

Products were from Sigma except when indicated. SOC medium was from Invitrogen, LB broth medium from Roth, ampicillin and Triton X100 from Euromedex, anti-protease tablets from Roche, Ni^2+^-NTA resin from Generon, DDM and DM from Anatrace, Amicon Ultra-15 devices from Millipore and Superdex 200 10/300 GL from GE.

##### BmrA expression

BmrA expression was adapted from methods previously reported (*23, 39*). The E504A mutant was generated and fused to a 6-histidine *N*-terminal Nickel-affinity tag in the pET15(+) plasmid and overexpressed in the CD43(DE3)Δ*acrB E. coli* strain, a gift of Pr. Klaas Martinus Pos. A freshly transformed colony was incubated in 3 mL LB containing 50 µg/mL for 7-8 h at 37 °C. Thirty microliters of this preculture were added to 1 L LB containing 50 µg/mL of ampicillin, which was then incubated at 22 °C until reaching 0.6 OD_600_. BmrA expression was induced by 0.7 mM IPTG followed by a 5-6 h incubation at 22 °C. Bacteria were collected at 5000 xg for 15 min., 4 °C and then suspended in 10 mL 50 mM Tris-HCl pH 8.0, 5 mM MgCl_2_ and 1 mM PMSF. Bacteria were lysed by 3 passages at 18,000 psi through a microfluidizer 100 (Microfluidics IDEX Corp). The solution was centrifuged 30 min. at 15,000 xg at 4 °C. The membrane fraction was pelleted by centrifugation for 1 h at 180,000 xg at 4°C, suspended in 50 mM Tris-HCl pH 8.0, 1 mM PMSF and 1 mM EDTA and centrifuged again. The final pellet was suspended in 20 mM Tris-HCl pH 8.0, 0.3 M sucrose and 1 mM EDTA, frozen in liquid nitrogen and stored at -80 °C.

##### BmrA purification

Membranes were solubilized at 5 mg/mL in 20 mM Tris-HCl pH 8.0, 100 mM NaCl, 15% glycerol (v/v), anti-protease tablets, 0.1 mM TCEP and 4.5% (w/v) Triton X100, under gentle agitation for 90 min. and then centrifuged 40 min. at 100,000 xg. The supernatant was loaded onto a Ni^2+^-NTA equilibrated in 20 mM Tris-HCl pH 8.0, 100 mM NaCl, 15% (v/v) glycerol, anti-protease tablets, 4.5% Triton X100 and 20 mM imidazole. The resin was washed with 20 mM Hepes-HCl pH 8.0, 100 mM NaCl, 20 mM imidazole, 1.3 mM DDM and 1 mM sodium cholate. The protein was eluted by adding 200 mM imidazole to the same buffer. BmrA fractions were pooled and diluted ten times in the Hepes buffer (same composition as above) without imidazole and loaded again on the same resin for another step of affinity chromatography.

The pool of BmrA fractions was concentrated on 50 kDa cutoff Amicon Ultra-15 devices, with the centrifuge speed set at 1000 xg for 10-15 min, and then loaded onto Superdex 200 10/300 using as mobile phase 20 mM Hepes-HCl pH 7.5, 100 mM NaCl, 0.7 mM DDM and 0.7 mM sodium cholate (DDM:cholate molar ratio of 1:1). The same step was also carried out at DDM- cholate ratio of 3:1 or 1:3. Cholate was systematically removed from the Superdex resin by a washing with 1M NaOH. The elution peak was then pooled and stored at 4 °C before use. BmrA was particularly stable when not concentrated as previously reported (*30*).

**Thermostabilisation assays** were carried out as previously reported (*30*). Membranes of BmrA diluted at 2 g proteins/L were solubilized with 10 mM DDM, with or without 1 mM of compounds **3a**-**3e** in a final volume of 2 mL, for 2 h at 4 °C. Solutions were clarified by centrifugation at 100,000 xg for 1 h at 4 °C and supernatants were aliquoted (50 μl) and individually submitted 30 min to a temperature of 25 to 90 °C using a PCR thermal cycler (PeqSTAR 2x gradient; Peqlab). Samples were then centrifuged 40 min at 20,000 xg and supernatants were analyzed by SDS-PAGE and Western-blot using anti-His antibody. The relative intensity of BmrA at each temperature was quantified on Western blot using Image Lab software 4.1 (Bio-Rad). Each condition was performed twice or thrice. Intensity was plotted as a function of the temperature and normalized. Data were fitted with equation 5 (see data fit section).

##### Detergents quantification

DDM bound to BmrA was quantified by mass spectrometry as described (*29*). Cholate was quantified as previously reported (*40*). Modelling of the detergent belt radius was done following the same protocol and using the DeltBelt server (www.deltbelt.ibcp.fr).

##### ATPase activity

The ATPase activity of BmrA was measured as previously described (*30, 41*). The protein in solution in 20 mM Hepes-HCl pH 7.5, 100 mM NaCl, 0.7 mM DDM and 0.7 mM cholate was diluted in the ATPase activity assay buffer containing either 1 mM DDM or a mixture of 0.7 mM DDM and 0.7 mM cholate, and the ATPase activity recorded.

##### Membrane-scaffold protein (MSP) production and purification

The MSP1E3D1 protein was expressed in BL21 *E. coli* (p1E3D1 plasmid, Addgene) as previously described (*32*). Bacteria were suspended in 50 mL of 40 mM Tris-HCl pH 7.4, 100 mM NaCl, 1 % (w/v) Triton X100, 0.5 mM EDTA, 1 mM PMSF. Two microliters of Benzonase (24 U/mL, Merck) were added and the bacteria were lysed by 2 passages at 18,000 psi through a microfluidizer 100 (Microfluidics IDEX Corp) and then centrifuged during 30 min. at 30,000 xg, 4°C. The supernatant was loaded onto a 0.5-mL Ni^2+^-NTA column (GE Healthcare) resin pre-equilibrated with 5 resin-volumes of 40 mM Tris-HCl pH 7.4, 100 mM NaCl, 1 % (w/v) Triton X100, 0.5 mM EDTA and 1 mM PMSF. The resin was then washed with 10 resin-volume with 3 different buffers: wash buffer 1 composed of 40 mM Tris-HCl pH 8.0, 300 mM NaCl and 1% (w/v) Triton X100; wash buffer 2 composed of 40 mM Tris-HCl pH 8.0, 300 mM NaCl, 50 mM sodium cholate and 20 mM Imidazole; wash buffer 3 composed of 40 mM Tris-HCl pH 8.0, 300 mM NaCl, 50 mM Imidazole. MSP1E3D1 was eluted with 15 mL of 40 mM Tris-HCl pH 8.0, 300 mM NaCl and 500 mM Imidazole. The factions of the elution pic were pooled and the TEV (2 mg/mL) was added to remove the His tag, at a ratio of 1 mg TEV for 40 mg MSP1E3D1. The mixture was then dialyzed (cutoff 12-14 kDa), a first time against 300 mL 40 mM Tris-HCl, pH 7.4, 100 mM NaCl and 0.5 mM EDTA for 3 hours and then against 700 mL of the same buffer, overnight at 4°C. After dialysis 20 mM imidazole was added and the solution loaded on a 0.5 mL Ni^2+^-NTA column equilibrated with 20 mM Tris-HCl pH 7.4 and 100 mM NaCl. The flow-through containing MSP1E3D1 was collected. The uncleaved fraction was eluted with 40 mM Tris-HCl pH 8.0, 300 mM NaCl and 500 mM Imidazole, dialyzed two times as above and finally concentrated spinning at 5,000 xg with a 100 kDa cutoff Amicon Ultra-15. The concentrated samples were frozen in liquid nitrogen and stored at -80°C.

#### BmrA nanodisc reconstitution

BmrA was reconstituted into nanodiscs as previously described (*42*) with the following modifications. Six hundred micrograms of purified BmrAE504A in 200 µL of Hepes-HCl pH 7.5, 100 mM NaCl, 0.035% DDM and 0.03% sodium cholate were mixed with 1.4 mg of *E. coli* lipids (Avanti Polar) in 56 µL of 99 mM cholate, 20 mM Hepes-HCl pH 7.5, 100 mM NaCl for 10 min at room temperature. The mix was then added of 665 µg MSP1E3D1 in 35 µL of 40 mM Tris-HCl, pH 7.4, 100 mM NaCl and 0.5 mM EDTA. The volume was completed to 1 mL with Hepes-HCl pH 7.5, 100 mM NaCl and incubated 1 h at room temperature. The final molar ratio of BmrA/MSP/lipids was 1/5/400 in 20 mM Hepes-HCl pH 7.5, 100 mM NaCl. SM-2 biobeads (170 mg/100 µg BmrA, Biorad) were then added to the mixture, placed 3 h under gentle agitation at room temperature. Empty nanodiscs were removed from the BmrA-nanodiscs by Ni^2+^-NTA chromatography. The resin was equilibrated with 20 mM Hepes-HCl pH 7.5, 100 mM NaCl, then loaded with the sample, washed with 20 mM Hepes-HCl pH 7.5, 100 mM NaCl, 20 mM imidazole. The BmrA-nanodiscs complex was then eluted with 20 mM Hepes-HCl pH 7.5, 100 mM NaCl, 200 mM imidazole. Imidazole was then removed from the solution by passing through a HiTrap desalting column equilibrated with 20 mM Hepes-HCl pH 7.5, 100 mM NaCl.

#### Ligand binding on BmrA in detergents

R6G, ATP-Mg^2+^ and cholate binding was carried out by incubating 15 min on ice 0.5 µM BmrA, WT or E504A mutant prepared in DDM or DDM-cholate with or without 5 mM ATP-Mg^2+^ in 20 mM Hepes-HCl pH 7.5, 100 mM NaCl, including 0.7 mM DDM and/or 0.7 mM cholate depending on the experiments. The binding of R6G was probed by intrinsic fluorescence recorded on a SAFAS Xenius spectrophotofluorimeter set up at a constant photo multiplicator voltage of 570 V. Tryptophan residues or *N*-acetyl tryptophan amide (NATA) used as negative control were excited at 280 nm, and their fluorescence emission spectra were recorded between 310 and 380 nm, with a 5-nm bandwidth for excitation and emission. Experiments were done in a quartz cuvette in a final volume of 200 µL, in which increasing amounts of R6G were added. Resulting emission curves were integrated and deduced from the same experiments carried out with NATA, used at the same concentration than that of BmrA tryptophan residues. Data were plotted as a function of R6G concentration. Binding of ATP-Mg^2+^ was carried out in the same way, pre-incubating BmrA E504A with or without 100 µM R6G for 15 min on ice.

#### Ligand binding to BmrA-nanodiscs complexes and empty nanodiscs

R6G, DDM and DM binding assays were carried out as above. Assays with empty nanodiscs (without BmrA) were carried out at the same nanodiscs concentration as that of BmrA-nanodiscs, complexes. This allowed to correct the fluorescence quenching due to the interaction between empty nanodiscs alone and ligands. Two cuvettes of NATA were also used: one for BmrA-nanodiscs complex and the other one for the empty nanodiscs. Data were analyzed in the same way as above.

**Doxorubicin transport by BmrA** was recorded as previously described (*23*). Ten micromolar of doxorubicin and 2 mM ATP were added to 100 μg *E. coli* inverted membrane vesicles containing overexpressed BmrA. Transport was initiated upon addition of 2 mM MgCl_2_ and monitored at 25 °C in 1-mL quartz cuvettes recording the fluorescence on a Photon Technology International fluorimeter at 590 nm with a bandwidth of 4 nm upon excitation at 480 nm with a bandwidth of 2 nm. Transport was initiated by adding 2 mM ATP-Mg^2+^. **R6G accumulation in *Bacillus subtilis* strains.** R6G accumulation assay was performed in *B. subtilis* strain 168 (WT) and 8R (overexpressing *BmrA* (*24*) kindly provided by Pr. Hans Krügel). Strains were grown overnight in LB medium at 37 °C with agitation, and then diluted to 0.05 OD_600nm_ with fresh medium. Once the culture reached 0.5 OD_600nm_, they were incubated with 5 µM R6G for 30 min more. Then 2 mL of each culture (∼ 1 OD_600nm_) was centrifuged at 15,000 xg for 10 min at 4 °C. The pellets were washed with 1 mL LB medium and centrifuged. The pellets were suspended in 500 µL of 50 mM Tris-HCl pH 8.0, 150 mM NaCl, 1 mg/mL lysozyme and incubated for 1 h at 37 °C with agitation. The cells were then incubated with 0.5% SDS for 15 min more. R6G fluorescence was recorded with a SAFAS Xenius spectrophotofluorimeter in a black 96 well-plate using 200 µL of cell lysates setting excitation to 526 nm and recording fluorescence between 541 and 650 nm. **Data fit**. Data were fitted using Microsoft Excel (365), SigmaPlot (v12.5) and GraphPad (v8) using/setting up the following equations:

Equation 1 (Intrinsic fluorescence quenching, ligand binding, one site saturation): f = *F_max_*\*abs([L])/(*K_D_*+abs([L])), *F_max_* = maximal intrinsic fluorescence without ligand, [L] = ligand concentration, *K_D_*, ligand dissociation constant.

Equation 2 (allosteric intrinsic fluorescence increase): f = *F_min_*+(*F_max_*-*F_min_*)/(1+([L]/*K_D_*)^-^*^h^*), *F_min_* = minimal intrinsic fluorescence without ligand, *F_max_* = maximal intrinsic fluorescence with ligand, [L] = ligand concentration, *K_D_*, ligand dissociation constant, *h* = Hill number.

Equation 3 (Sigmoidal, 3 parameters): f = *F_max_*/(1+exp(-([L]-[L]*_50_*)/*b*)), *F_max_* = maximal intrinsic fluorescence, [L] = ligand concentration, µM, [L]_50_ = ligand concentration at half-maximal intrinsic fluorescence, µM.

Equation 4 (Intrinsic fluorescence quenching, ligand binding, two sites saturation): f = *F_max1_*\*abs([L])/(*K_D1_*+abs([L])) + *F_max2_*\*abs([L])/(*K_D2_* + abs([L])), *F_max1,_ F_max2_* = maximal intrinsic fluorescence without ligand, [L] = ligand concentration, *K_D1_*, *K_D2_*, ligand dissociation constants.

#### HDX experiments

HDX-MS experiments were performed using a Synapt G2-Si mass spectrometer coupled to a NanoAcquity UPLC M-Class System with HDX Technology (Waters^™^). All the reactions were carried out manually. Labeling was initiated by diluting 5 µL of typically 15 µM BmrA in nanodiscs, in 95 µL D_2_O labeling buffer, 5 mM Hepes pD 8.0, 50 mM NaCl. For the Vanadate-trapped condition, the labeling buffer additionally contained 10 mM ATP, 10 mM MgCl_2_ and 1 mM Vanadate. Samples were labeled for 2, 5, 15 and 30 minutes at 20 °C. Subsequently, the reactions were quenched by adding 22 µL of ice-cold quenching buffer, 0.5 M glycine, 8 M guanidine-HCl pH 2.2, 0.035% DDM and 0.03% sodium cholate, to 100 µL of labelled sample, in ice bath. After 1 min, the 122-µL quenched sample was added into a microtube containing 200 µg of activated zirconium magnetic beads (MagReSyn Zr-IMAC from Resyn Biosciences, USA), to remove the phospholipids (*43*). After 1 min. magnetic beads were removed and the supernatant injected immediately through a 100-µL loop. Labelled proteins were then subjected to in-line digestion at 15 °C using a pepsin column (Waters Enzymate™ BEH Pepsin Column 300 Å, 5 µm, 2.1 x 30 mm). The resulting peptides were trapped and desalted for three minutes on a C4 pre-column (Waters ACQUITY UPLC Protein BEH C4 VanGuard pre-column 300 Å, 1.7 µm, 2.1 x 5 mm, 10K - 500K) before separating them with a C4 column (Waters ACQUITY UPLC Protein BEH C4 Column 300 Å, 1.7 µm, 1 x 100 mm) using a linear acetonitrile gradient of 5-40% in 15 min and then four alternative cycles of 5% and 95% until 25 min.. The valve position was adjusted to divert the sample after 11.2 min of each run from C4 column to waste to avoid contaminating the mass spectrometer with detergent. Two full kinetics were run for each condition, one after the other, to get duplicate of each deuteration timepoint. Blanks, with equilibration buffer, 5 mM Hepes pH 8.0, 50 mM NaCl, were injected after each sample injection and pepsin column washed during each run with pepsin wash (1.5 M guanidine-HCl, 4% acetonitrile, 0.8% formic acid pH 2.5) to minimize the carryover. Electrospray ionization Mass spectra were acquired in positive mode in the m/z range of 50−2000 and with a scan time of 0.3 s.

For the identification of non-deuterated peptides, data was collected in MS^E^ mode and the resulting peptides were identified using PLGS™ software (ProteinLynx Global SERVER 3.0.2 from Waters™). Deuterated peptides were identified using DynamX 3.0 software (Waters™), using the following parameters: minimum intensity of 1000, minimum products per amino acid of 0.3 and file threshold of 2. Deuteros 2.0 software (*44*) was used for data analysis, visualization and statistical treatments. The mass spectrometry data have been deposited to the ProteomeXchange Consortium via the PRIDE (*45*) partner repository with the dataset identifier PXD022185.

#### Biophysics

##### Products

Crystallization solutions were from Grenier bio-one. The Mosquito crystallization robot is from TTP Labtech. Crystallization plates and cover were from Grace Bio-Labs. The cryoprotection kit was from Molecular Dimensions. Cryschem plates were from Hampton Research. Vitrobot grid freezing device is from FEI. The Talos Arctica and Titan Krios G3 are from Thermo Scientific.

##### X-ray

###### Protein crystallization

The crystallography step was performed at 19 °C. Crystals were obtained by vapor diffusion on hanging drops. E504A BmrA mutant was concentrated by centrifugation-filtration to 7-10 mg/mL spinning at 500 xg on a 50 kDa cutoff Amicon Ultra-15 at 22 °C. BmrA E504A mutant was then incubated with 5 mM ATP-Mg for 30 min. Crystallogenesis was done by mixing with a Mosquito 450 nL of reservoir solution containing 100 µL 0.1 M Tris-HCl pH 8.5, 23-27% PEG 1000 with 50 nL of compounds 3a-3e and 500 nL of BmrA E504A sample. the mix was deposited on a plastic cover, sealed onto the plate and imaged periodically with a Formulatrix. Crystals appear after 3 days, grown up to 5-8 days and progressively disappeared if the incubation lasted longer.

###### Crystal cryocooling

As BmrA E504A mutant crystals were sensitive to cryoprotection, it was therefore performed using the CryoProtX MD1-61 kit. Best results were obtained with a final solution containing 12.5% (v/v) di-ethylene glycol, 18% (v/v) 2-methyl-2,4-pentanediol, 7% (v/v) ethylene glycol, 12.5% (v/v) 1,2-propanediol, 12.5% (v/v) dimethyl sulfoxide, supplemented with 5 mM ATP-Mg and 1 mM compounds **3a-3e**. One microliter of cryo-solution was divided in 3 drops under the binocular, close to the drop containing the crystal and then gently brought in contact using the freezing loop, at the opposite side where the crystal was sitting, in the course of 1 min. This operation was performed in Cryschem sitting drop plates, with the drop sitting in the middle of a water-filled reservoir to saturate the solution with humidity. Crystals were then harvested and placed on a fresh drop of cryo-solution for 1 min. before harvesting and cryocooling in liquid N_2_. Crystals were stored in liquid N_2_ before being analyzed at the synchrotron.

###### Diffraction data acquisition

Diffraction screening has been performed at ESRF and SOLEIL synchrotrons on multiple beamlines over the years. Best data set was collected on PX2 at SOLEIL, consisting of a low-resolution pass at low transmission, and a high-resolution dataset at full transmission collected helicoidally. Crystal polymorphism was strongly present, precluding data merging among several crystals. Crystals diffracted very anisotropically, going to 3.9 Å resolution in the strongest diffracting direction. Data were processed in XDS as spherical to the highest resolution possible (3.9 Å) even though spherical statistics were not usable. Staraniso analysis for diffraction anisotropy revealed that completeness was 78% in the highest resolution shell, therefore revealing that all the data collectable for this crystal had been collected. Data was cut at the diffraction limits suggested by the Staraniso server. Anisotropic diffraction table is available in supplementary Table 1.

###### X-ray structure and model building

Phases were solved by molecular replacement using Phaser on amplitudes, with data corrected for anisotropy using Staraniso, and using the outward facing conformation dimer of Sav1866 (PDB code ID 2hyd) or MsbA (PDB code ID 3b60) as search models. Although Sav1866 and MsbA structures are very similar, the MsbA model yielded higher molecular replacement solution scores. Crystals belonged to the P21 space group with 2 dimers in the asymmetric unit. The molecular replacement solution was clear, but the electron density was very noisy due to the large conformational changes observed on BmrA, and that resulted in poor overall phases. The core of the protein was nevertheless clearly visible with helices as tubes. The nucleotide-binding domain was very blurry as well as external loops. The model was turned into poly-Ala to place helices of the transmembrane region, and initial movement of the TM1-TM2 hinge. Refinement was carried out in autoBUSTER using corrected amplitudes, applying strict NCS. Iterative manual building in Coot followed by refinement resulted in visible continuous electron density with decreasing R-factors. Density for large amino acids appeared, as well as for ATP. Sequence was assigned, and iterative refinement continued with introduction of TLS refinement (1 TLS per chain, 4 total). It yielded R-factors around 30 and 35 for R and Rfree respectively, with small grooves in the helices. Re-definition of TLS (1 for a dimer of TMD, 1 for a dimer of NBD, 4 total) resulted in a dramatic decrease of R-factors by 3 points, and much clearer electron density features, helices with large grooves and side chain density. Unwinding of TM3 next to residue 136 was apparent, as well as helix breaks in the trans-membrane region and clear density for ATP. Some incorrect modeling of ATP became apparent with negative and positive density showing where the correct position was then defined. NCS was relaxed and correct modeling of geometry clashes was carried out in ISOLDE. Registry was built by starting to assign using initial first large density features clearly visible as refinement converged for the TMD, then using superpositions for the NBD. Registry at key locations was then probed by replacing several amino-acids, or by trying to turn helices by one amino-acid clockwise or counterclockwise and probed by refinement. Newly refined structures clearly showed positive or negative densities indicative of incorrectly modeled features. Ramachandran and rotamers outliers were corrected, yielding a final model with R = 26.0% and Rfree = 32.1%. The final model was deposited in the Protein Data Bank under the accession code 6r72.

##### Cryo-EM

###### Sample preparation

Purified BmrA E504A mutant at 3.4 mg/mLin DDM-cholate 1:1 (0.035%- 0.03%) was incubated with 0.1 mM R6G followed by 5 mM ATP-Mg. Three microliters of this mixture were applied to cryo-EM Au-grids (Cflat 1.2/1.3 3Au) previously discharged in air for 40 s at 20 mA (PELCO easiGlow), blotted for 3 s, and plunge frozen in liquid ethane with a Vitrobot grid freezing device.

###### Data acquisition

Image processing. Best grids screened with a Talos Arctica were then imaged with a Titan Krios G3 electron microscope equipped with a K2 camera and operating at 300 keV. A total of 3477 movies of 40 frames each were acquired over 2 data-collection sessions in electron counting mode at 1.06 Å/pixel, 6.4 electrons/pixel/s, with a total exposure time of 6 s and combined into a single MRC stack using EPU automatic data collection control software using defocus values ranging from 1.2 to 3.2 µm. Contrast transfer function (CTF) parameters were estimated from the averaged movie with CTFFIND4 and 2170 particle images were selected manually and subjected to 2D classification in cryoSPARC v2. Automatic particle selection was performed with templates from the 2D classification. Beam induced particle motion between fractions was corrected with a new implementation of alignparts lmbfgs in cryoSPARC v2. The number of particle images were reduced to 128372 by further 2D and 3D classifications and refinements. Models were calculated ab initio and refined without the application of symmetry with cryoSPARC v2. For each data collection session automatically picked particles were cleaned with 2 rounds of 2D classification followed by a preliminary round of 3D classification to further remove obvious junk particles such as empty detergent micelles that were not eliminated during the 2D classification process. Although no discrete conformation could be isolated to high resolution, removal of additional particles improved the resolution of the final maps suggesting significant non-discrete or continuous flexing. Since these maps suggested a significant amount of small, non-discrete flexing, better resolved maps were obtained using cryoSPARC v2’s non-uniform refinement feature. An additional refinement with the application of C2 symmetry was performed that resulted in a gain of 0.3 Å in overall resolution which helped to slightly improve the interpretability of the map in the model building process. The asymmetric and C2 symmetrized maps have been deposited in the Electron Microscopy database under the accession codes EMD-4749 and EMD-12170 respectively.

###### Model building and refinement

The X-ray model was docked into a 3.9 Å C2 symmetrized cryo-EM density map and improved with iterative rounds of manual building in Coot and Isolde followed by real_space_refine in Phenix. Of Note, sharpening the C2 symmetry map using Phenix led to improved features in the trans-membrane domain, but worse in outer loops and the NBD. The final model was thus built using both sharpened and unsharpened maps. The final model was validated with MolProbity and EMringer and deposited in the Protein Data Bank under the accession code 6R81 and electron microscopy database EMDB-4749.

Two small densities were visible in the C2 symmetrized map at the locations of R6G. Re-examination of the data with no symmetry led to the identification of clearer densities in which R6G could be placed and suitably refined. Notably, both densities are not equivalent in the two halves of BmrA, suggesting that in both binding sites there is a heterogeneity/flexibility of binding, reminiscent of substrate release. Understandably, the application of C2 symmetry masked the quality of the reconstructions at these locations since these sites are not identical with respect to R6G binding. Thus BmrA was refined in the presence of R6G, following the same procedure as above using the asymmetric map. Final model and maps were deposited in the Protein Data Bank under the accession code 7BG4 and electron microscopy database EMD-1. 12170. Model statistics are provided in supplementary Table 1.

#### Bioinformatics

Both the X-ray and the cryo-EM(C2) structures span residues 10 to 589, and both miss a few residues (271-278 in the X-ray structure, 273-278 in the cryo-EM structure), corresponding to the loop region between TM5 and TM6. Complete models of dimeric wild-type BmrA were generated using Modeler (v9.12), for both the X-ray and the cryo-EM structures, using the structure of the ABC transporter related protein from *Novosphingobium aromaticivorans* (PDB code ID 4mrs) (*46*) as a template for the missing residues, and the alignment generated by HHPred (*47*). The N-termini were capped with acetyl groups, and the missing N-terminal residues were not modeled. Both models contained ATP molecules and Mg^2+^ ions, as observed in both the X-ray and the cryo-EM structures. The models were then oriented using the OPM server (http://sunshine.phar.umich.edu/server.php) (*48*) and embedded into a mixed POPE/POPG bilayer (ratio 3/1) using the CHARMM-GUI membrane builder (*49*), and the replacement method. The systems were solvated and 150 mM KCl was added to the solution, yielding a total of ∼157,000 atoms in tetragonal boxes of dimensions ∼100x100x165 Å^3^.

All simulations were run with the GROMACS (v2016.4) software package (*50, 51*). The CHARMM36 force field was used for both the lipids and the protein, together with the CHARMM TIP3P water model. Non-bonded interactions were calculated with a cutoff of 1.2 Å, with a shift function on the potential to avoid discontinuities. Neighbor lists were updated using the Verlet scheme. Long-range electrostatic interactions were calculated with the Particle Mesh Ewald method (*52*). Bonds involving hydrogen atoms were constrained using the P-LINCS algorithm (*53*).

Each system was minimized by steepest descent and then equilibrated using a 6-cycle equilibration scheme, using position restraints on the protein and gradually reducing the force constant. Equilibration and production runs were performed at 303.15 K and 1 bar; the temperature was kept constant with the velocity rescale algorithm (*54*) and the pressure with the Parrinello-Rahman barostat (*55*). The integration time step was set to 2 fs. For each system, four replicates were simulated for 500 ns each. The first 200 ns of each simulation were treated as equilibration, and average quantities (average structures, inter-atomic distances, RMSD, RMSF, B factors) were computed on the remaining 300 ns. Two additional replicates were run with long equilibration steps (275 ns before 500 ns of production) to confirm the closing movement of the cavity.

Due to the relatively large size and the transmembrane nature of BmrA, we expect functional motions of the transport cycle to take place on time scales much longer than the simulation time (probably 3-4 orders of magnitude longer), which are currently not accessible by all-atom molecular dynamics. Therefore, we only expect to observe relatively fast conformational changes, and changes driven by strong driving forces.

---

**Fig. S1.**
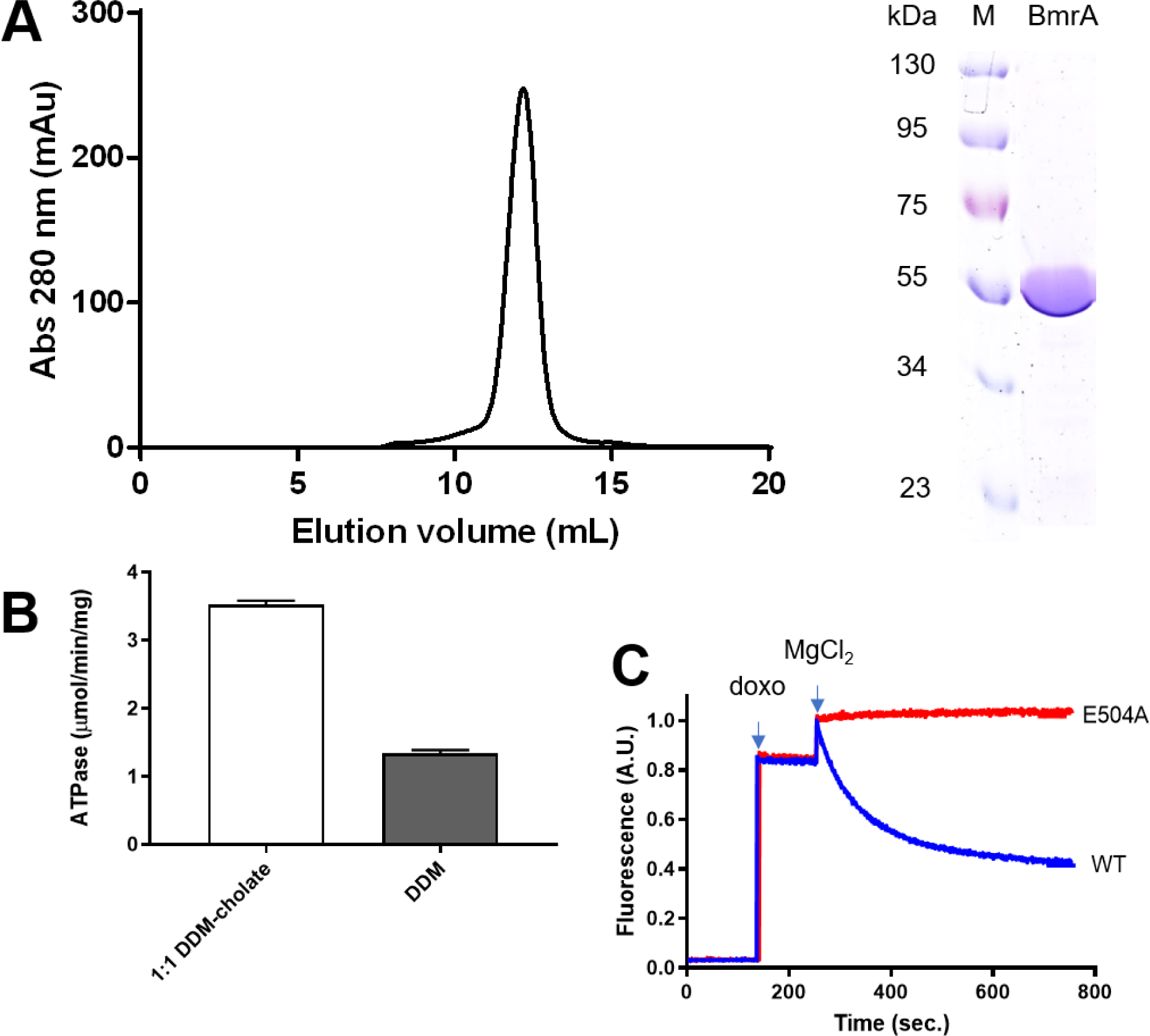
Purification of BmrA, ATPase activity and transport assay. **(A)** Preparative SEC profile of detergent purified BmrA (left panel). The peak fraction was analyzed by SDS-Page (right panel). **(B)** ATPase activities of WT BmrA purified with DDM or DDM-cholate mixture. **(C)** Doxorubicin (doxo) transport activity of WT BmrA and the inactive E504A mutant.

**Fig. S2.**
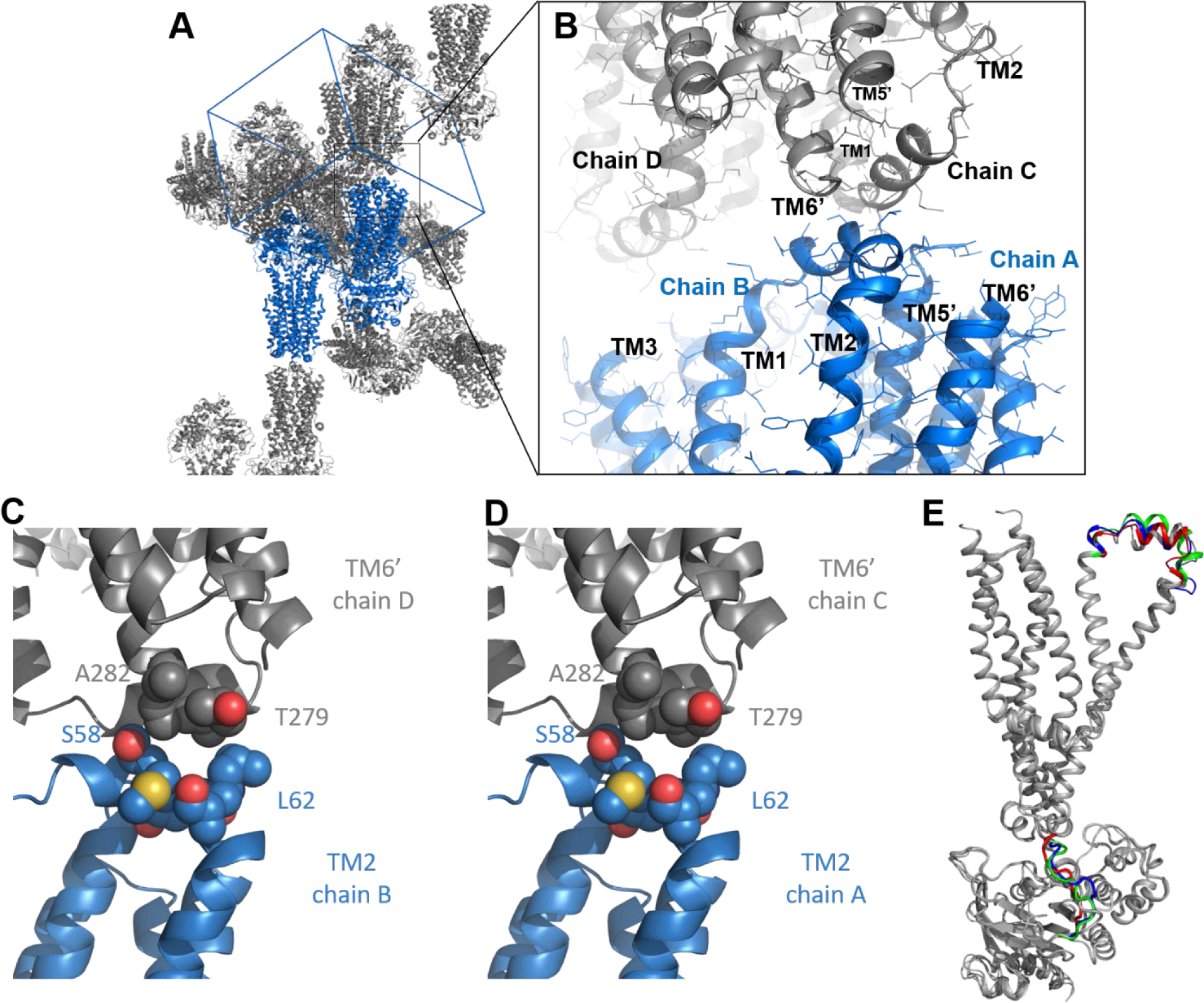
Crystallographic packing and difference between monomers in the asymmetric unit. **(A)** Overall crystal packing. The 4 BmrA monomers A-D of the asymmetric unit, assembled in 2 dimers, AB colored in blue, and the symmetric dimer CD in grey. Proteins are represented as cartoon, and the cell is drawn in blue. (**B**, **C, D**) Close-up views of the interaction between TM1-2 of the B/D and C/A monomers. **(E)** Differences between monomers in the X-ray structure. Structures are represented in cartoon, colored in grey. Flexible regions are highlighted in green (chain B), blue (chain C) and red (chain D). Each 4 monomers of the asymmetric unit of the crystallographic structure were superposed onto chain A.

**Fig. S3.**
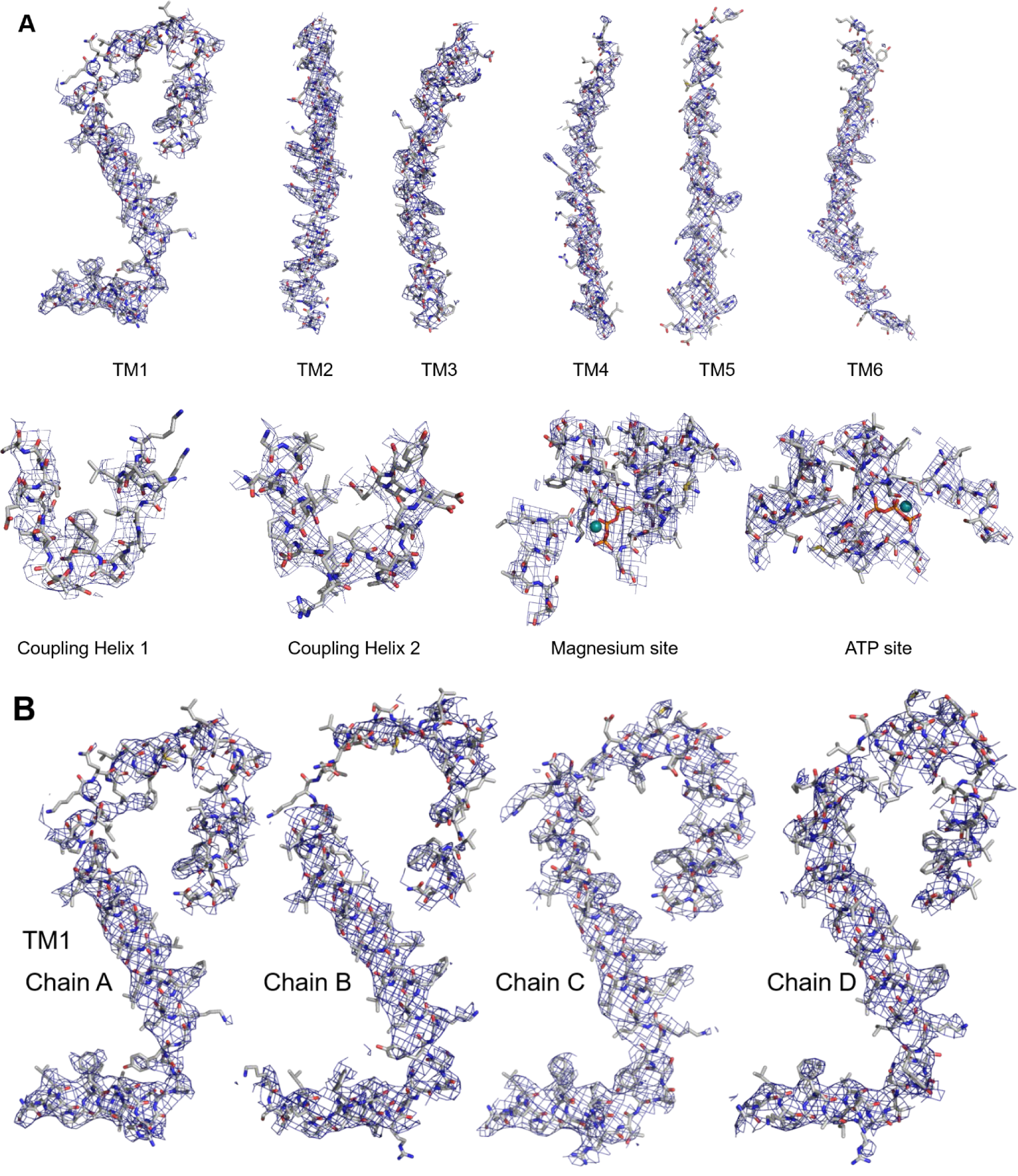
X-ray densities of BmrA. **(A)** TM helices, coupling helices and ATP-Mg^2+^ binding site of chain A. **(B)** Densities of the TM1-extracellular loop 1 for each chain.

**Fig. S4.**
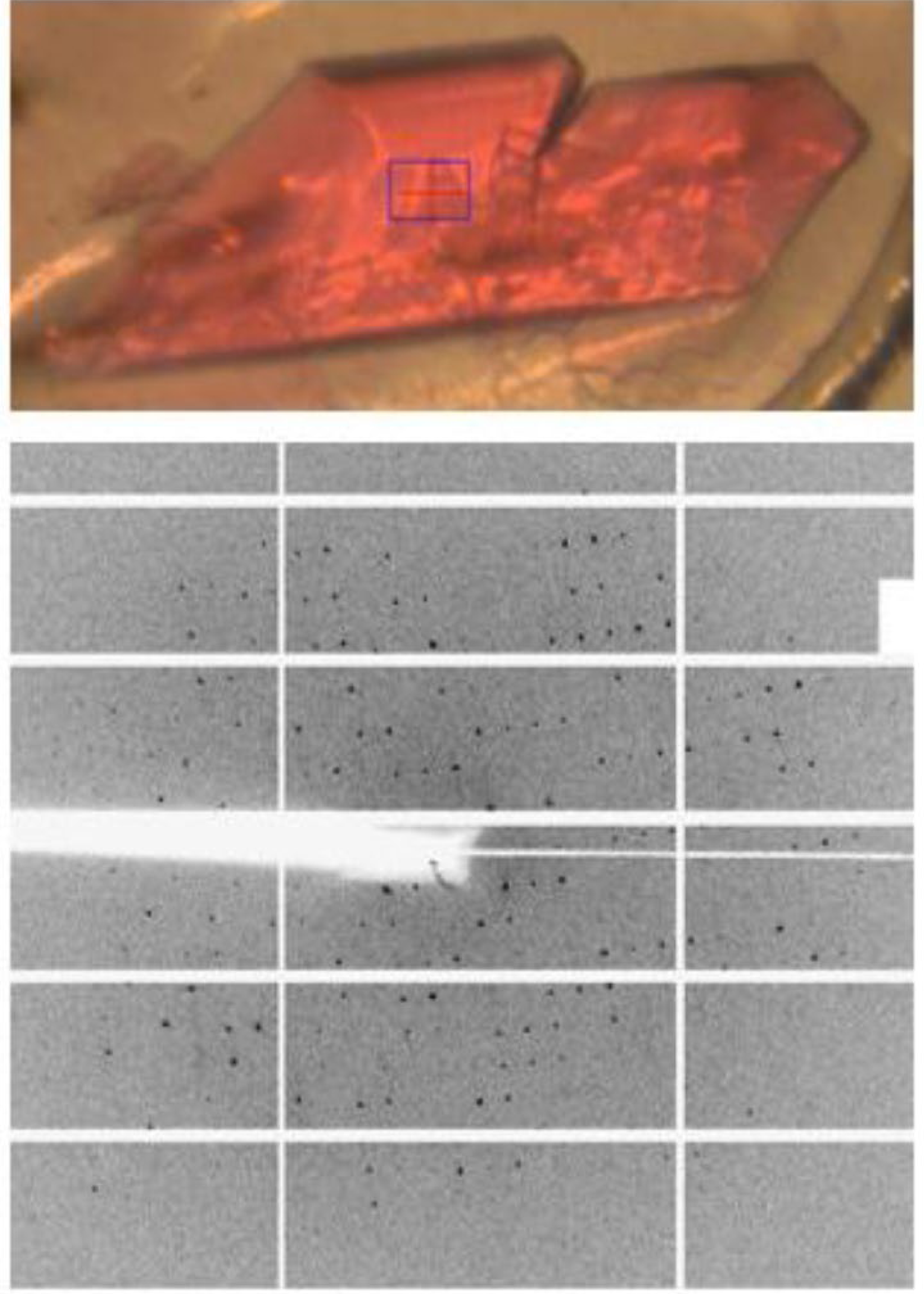
Crystallization of BmrA E504A in complex with ATP-Mg^2+^ and R6G.

**Fig. S5.**
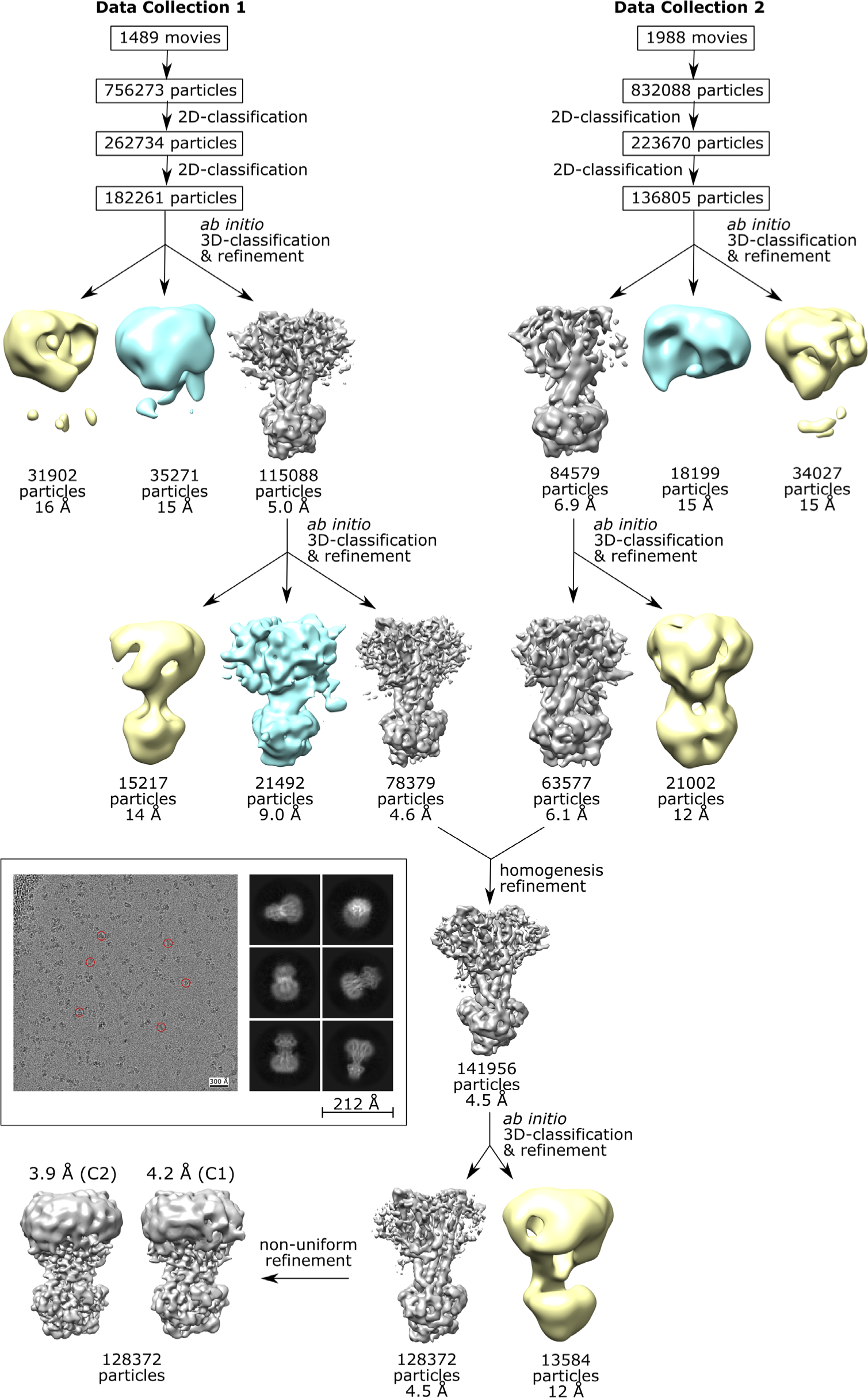
Image collection, 2D-3D classification, and processing workflow of cryo-EM image analysis of BmrA in OF conformation. Micrographs from two separate data collection sessions were processed in parallel and the best particles from each session were later combined to produce the final maps. For each session, picked particles were cleaned using 2 rounds of2D classification followed by a 3D classification. Heterogeneity within the dominating outward- facing conformation was further assessed with additional rounds of 3D classification that removed an additional 35 % of the outward-facing particles. Although removal of these particles improved the resolution of the dominating outward-facing conformation, discrete conformations could not be refined to high resolution suggesting that a large degree of small, non-discrete flexing was interfering with particle alignment. Due to the significant flexing amount, the final maps were refined using cryoSPARC’s non-uniform refinement feature resulting in better resolved maps. An additional refinement with the application of C2 symmetry was performed that resulted in an improvement in resolution. **Boxed**: example micrograph with particles used for 2D classification (red circles), and corresponding representative 2D class averages.

**Fig. S6.**
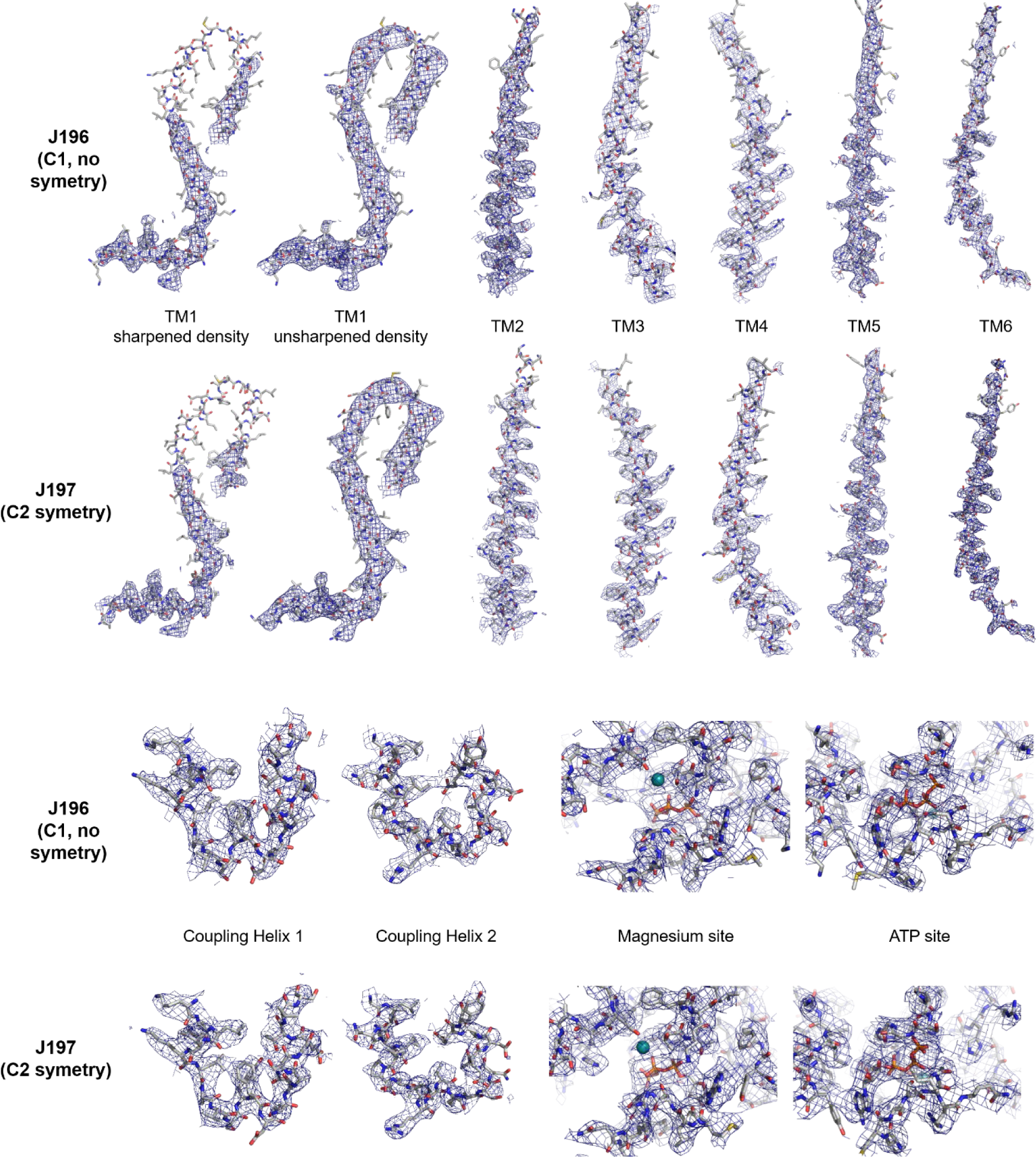
Cryo-EM densities of BmrA. Sharpened EM densities of the TM helices (and also unsharpened for TM1), coupling helices and the Magnesium and ATP binding sites for J196 (C1, no symmetry) and J197 (C2 symmetry map).

**Fig. S7.**
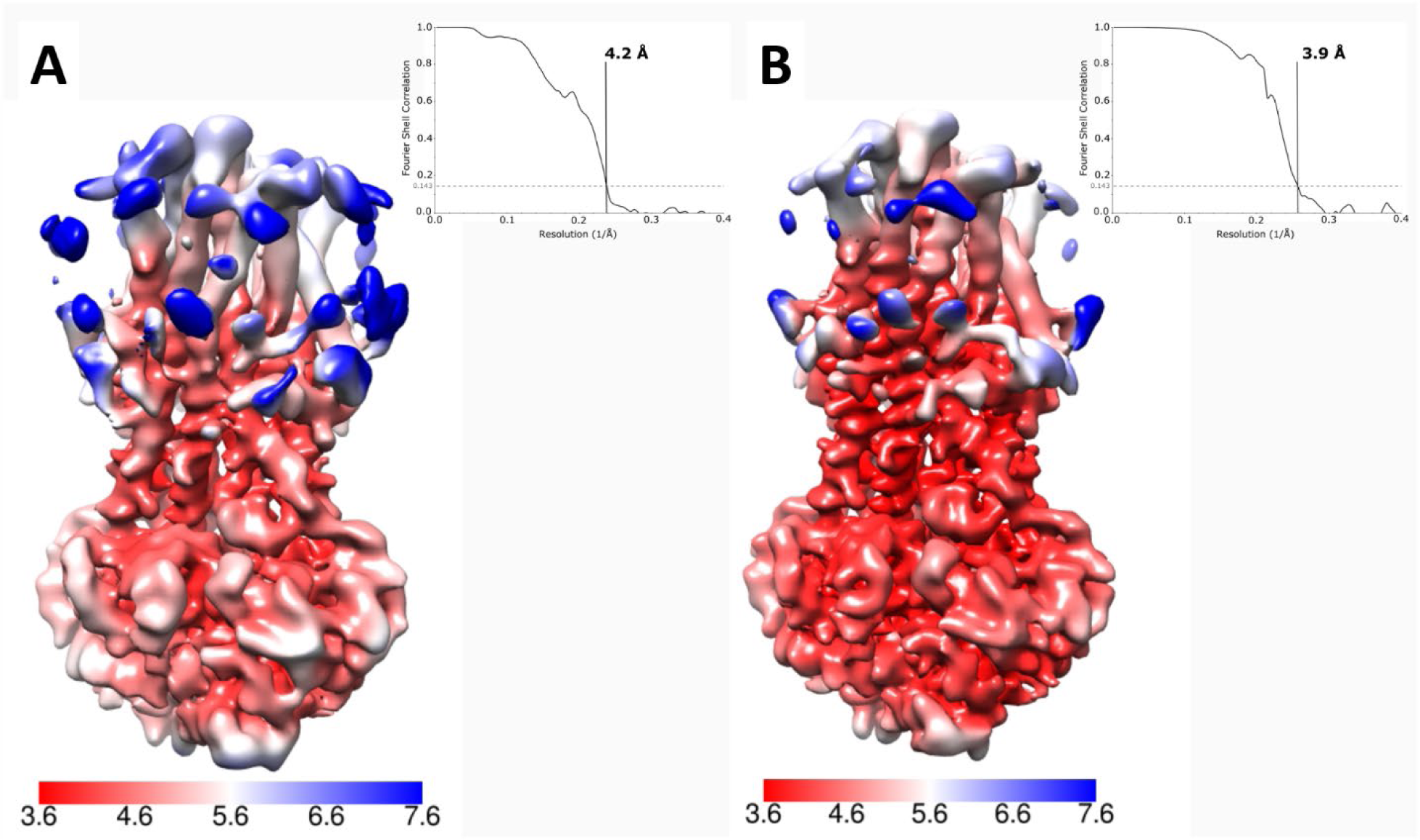
Assessment of the cryo-EM data. Local-resolution estimation of the C1 (A) and C2 (B) density maps and their corresponding Fourier Shell Correlation (FSC).

**Fig. S8.**
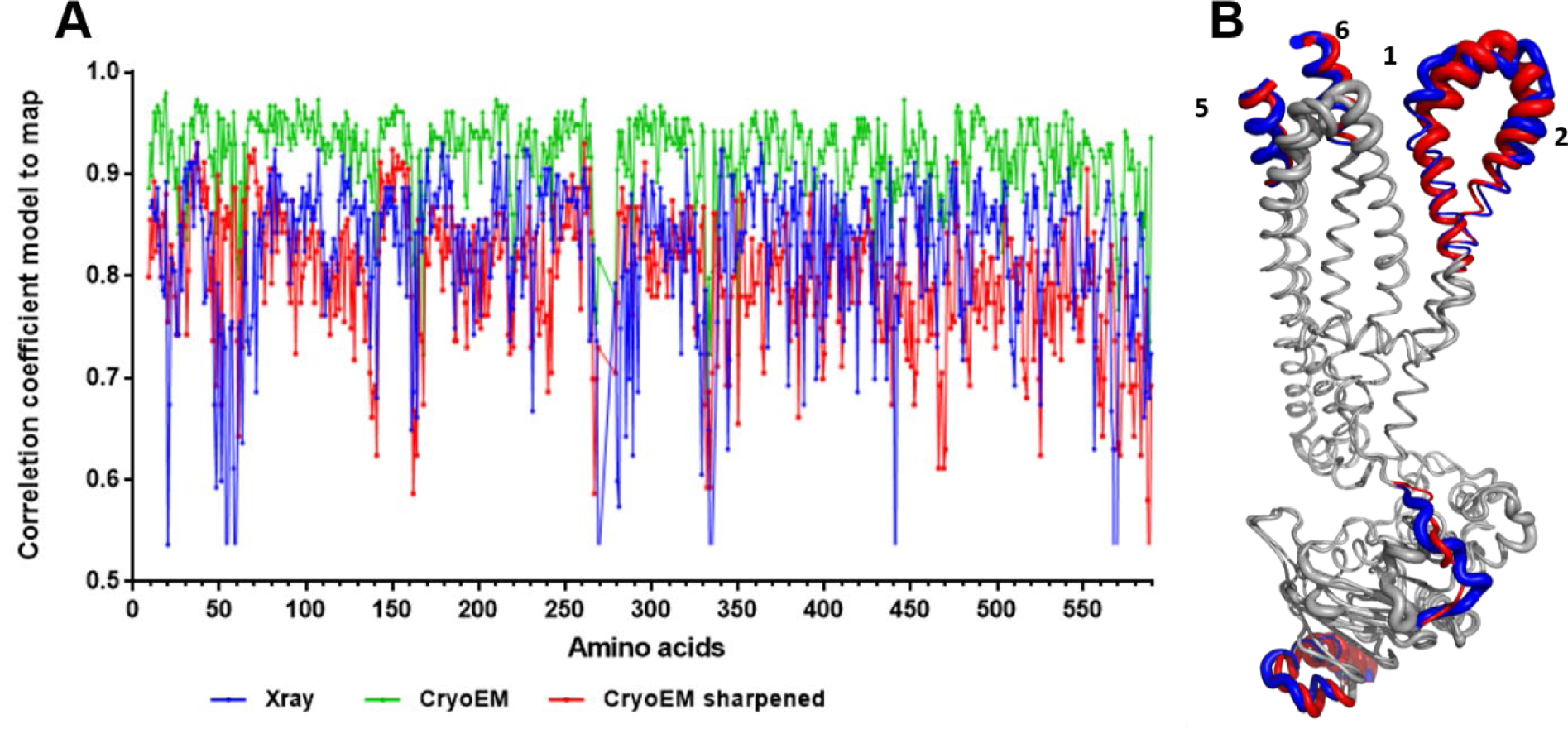
Correlation coefficient model to map and identification of the flexible parts of X-ray and cryo-EM structure. **(A)** The value of correlation coefficient model to map was calculated for each model against the corresponding density map. The results are plotted as a function of the amino acid sequence (X-ray in blue, cryo-EM unsharpened in green and sharpened in red). **(B)** The X-ray and the cryo-EM structures are superposed; the flexible parts are colored in red and blue for the cryo-EM and X-ray structure, respectively. These flexible parts correspond to the lowest CC values. The cartoon is represented with the thickness of the sausage corresponding to the B-factor, the higher the B-factor, the larger the sausage.

**Fig. S9.**
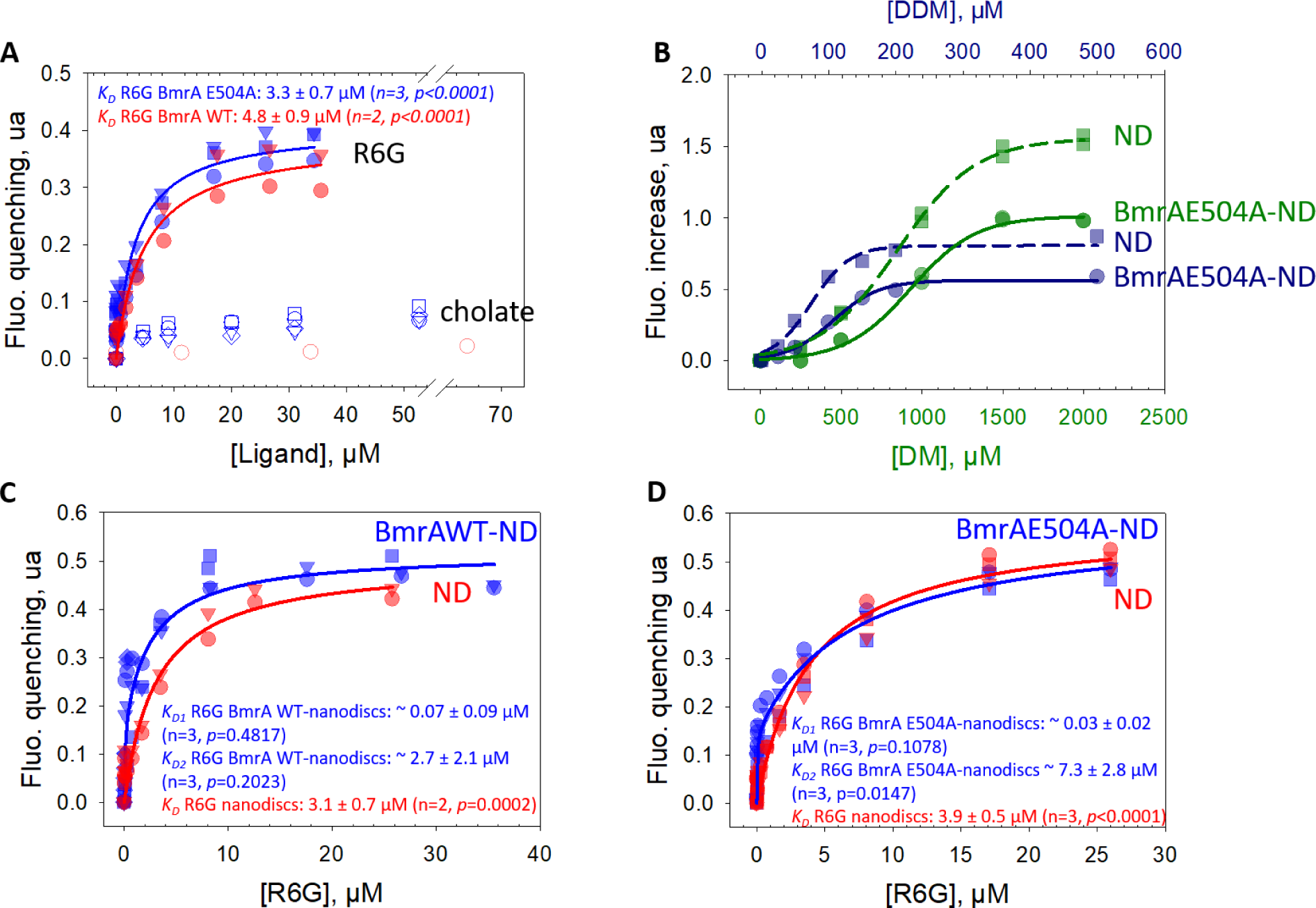
Binding of compounds to BmrA probed by intrinsic fluorescence. **(A)** Binding of R6G (filled symbols) and cholate (empty symbols) to BmrA WT (red) or E504A mutant (blue) purified in DDM. Data were fitted with equation 1. No significant fluorescence change was observed upon cholate addition in same conditions. **(B)** Effect of DDM (blue) and decyl maltoside (DM, green) on empty nanodiscs (ND) and BmrA-nanodiscs complexes. BmrA E504A was purified in DDM and then reconstituted into nanodiscs to which DM or DDM were added. The same experiments were done with empty nanodiscs (ND). Data were fitted using equation 3, giving a half-maximal fluorescence increase detergent concentration, [DDM]_50_, of ∼106 ± 6 µM (n = 1, *p* < 0.0001) and ∼74.1 ± 7.7 µM (n = 1, *p* < 0.0006), and [DM]_50_ of 931 ± 23 µM (n = 2, *p* < 0.0001) and 850 ± 21 µM (n = 2, *p* < 0.0001) for the BmrA-nanodiscs complexes and empty nanodiscs, respectively. **(C, D)** Binding of R6G to BmrA WT-nanodiscs (**C**, blue), BmrA E504A- nanodiscs (**D**, blue) and corresponding empty nanodiscs (**C or D**, red). The amount of empty nanodiscs used in these experiments correspond to that of MSP1E3D1 proteins in complex with BmrA, estimated by SDS-PAGE using each purified protein as standard. Data best fitted with equation 1 for empty nanodiscs (one site saturation) and equation 4 for BmrA-nanodiscs complexes (two sites saturation).

**Fig. S10.**
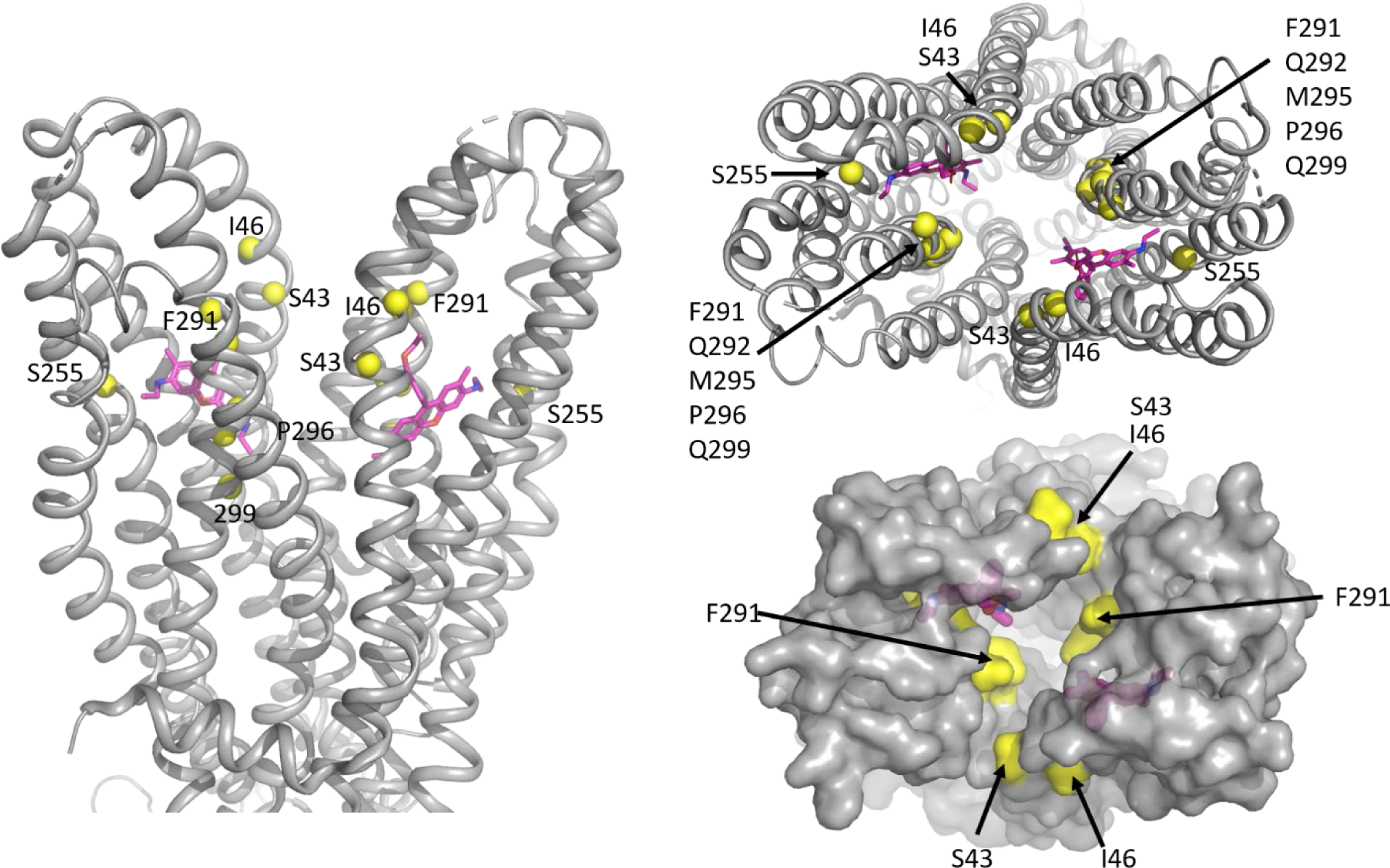
BmrA residues equivalent to those of the human ABCB1 involved in Taxol binding. Cryo-EM BmrA structure is displayed in grey in which residues in yellow correspond to those involved in taxol binding in the human ABCB1 (*14*).

**Fig. S11.**
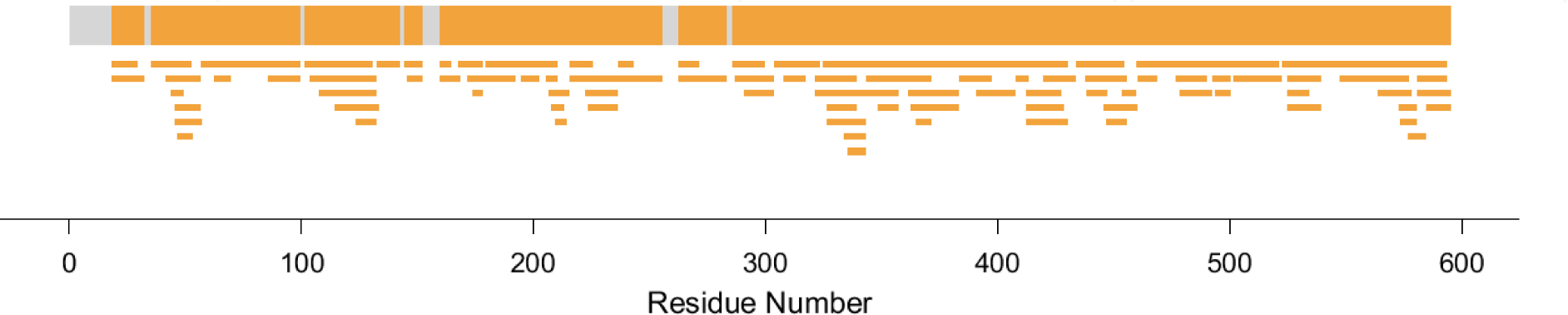
BmrA peptide coverage map obtained in the HDX-MS experiment. The common peptides identified in both states (apo and Vi-trapped) of WT BmrA reconstituted in nanodiscs are indicated in orange. The overall sequence coverage was approximatively 93%.

**Fig. S12.**
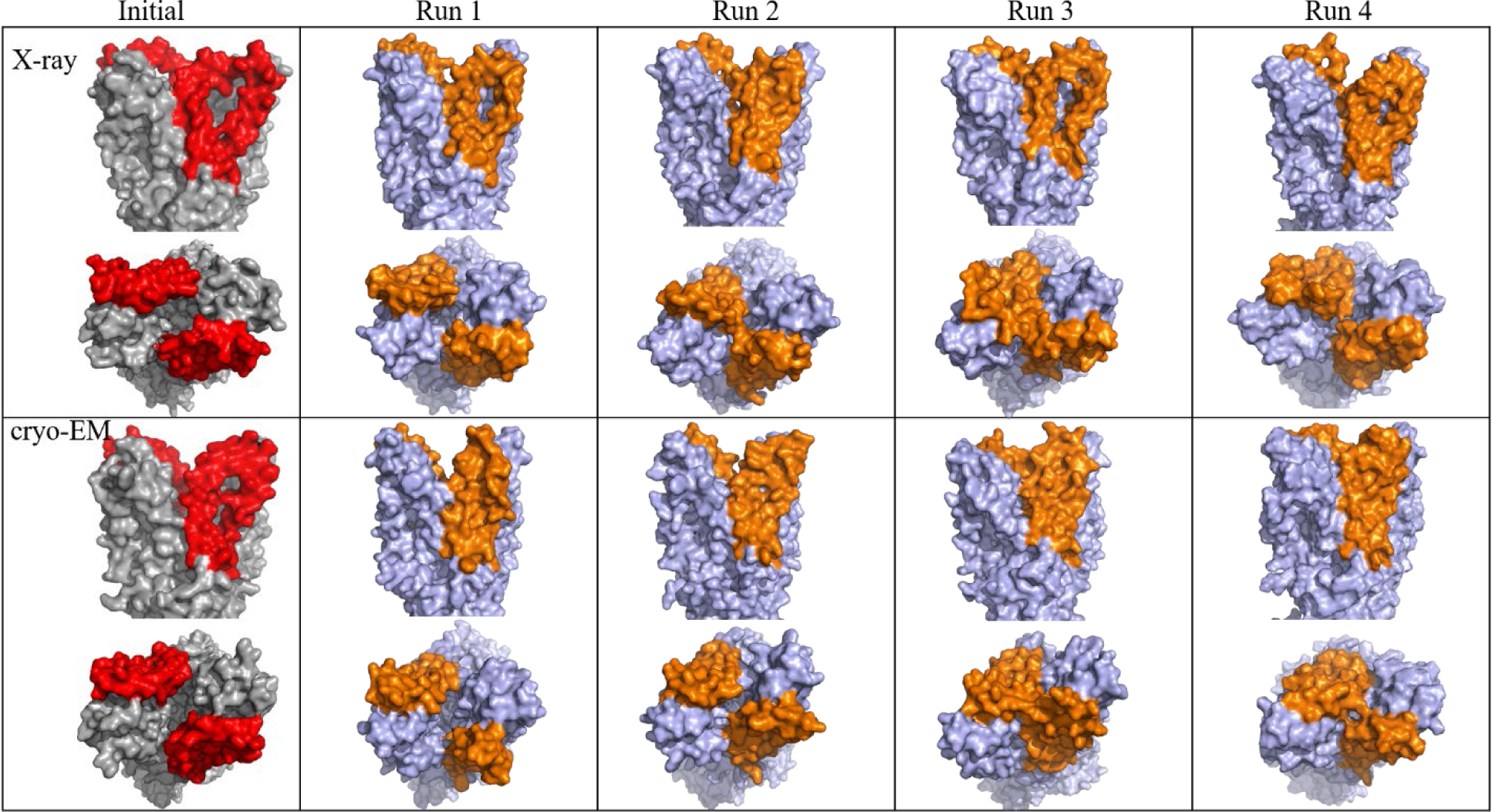
Molecular dynamic simulation results. The X-ray and cryo-EM structures are in grey and red (TM1-2). The final structures after 500 ns of simulation resulting from the four simulation runs are in light blue and orange (TM1-2). The side view (upper panel) and the top view (lower panel), from the outside of the membrane, are shown for each simulation.

**Fig. S13.**
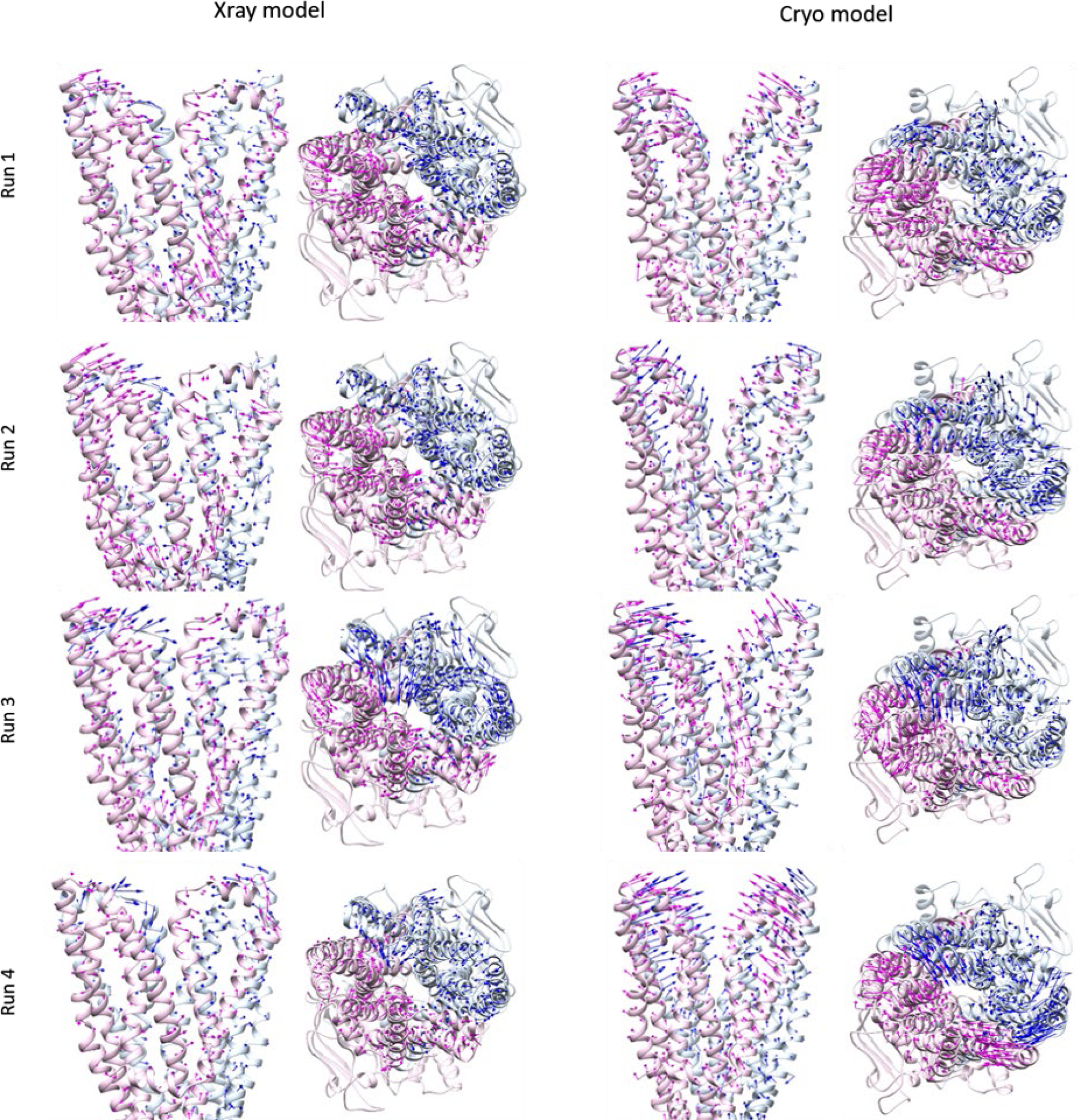
Conformational changes of the TM region in the 8 simulations. Chain A is shown in pink and chain B in blue. Arrows indicate the displacement of the TM regions from the starting structures to the average structures.

**Table S1:**
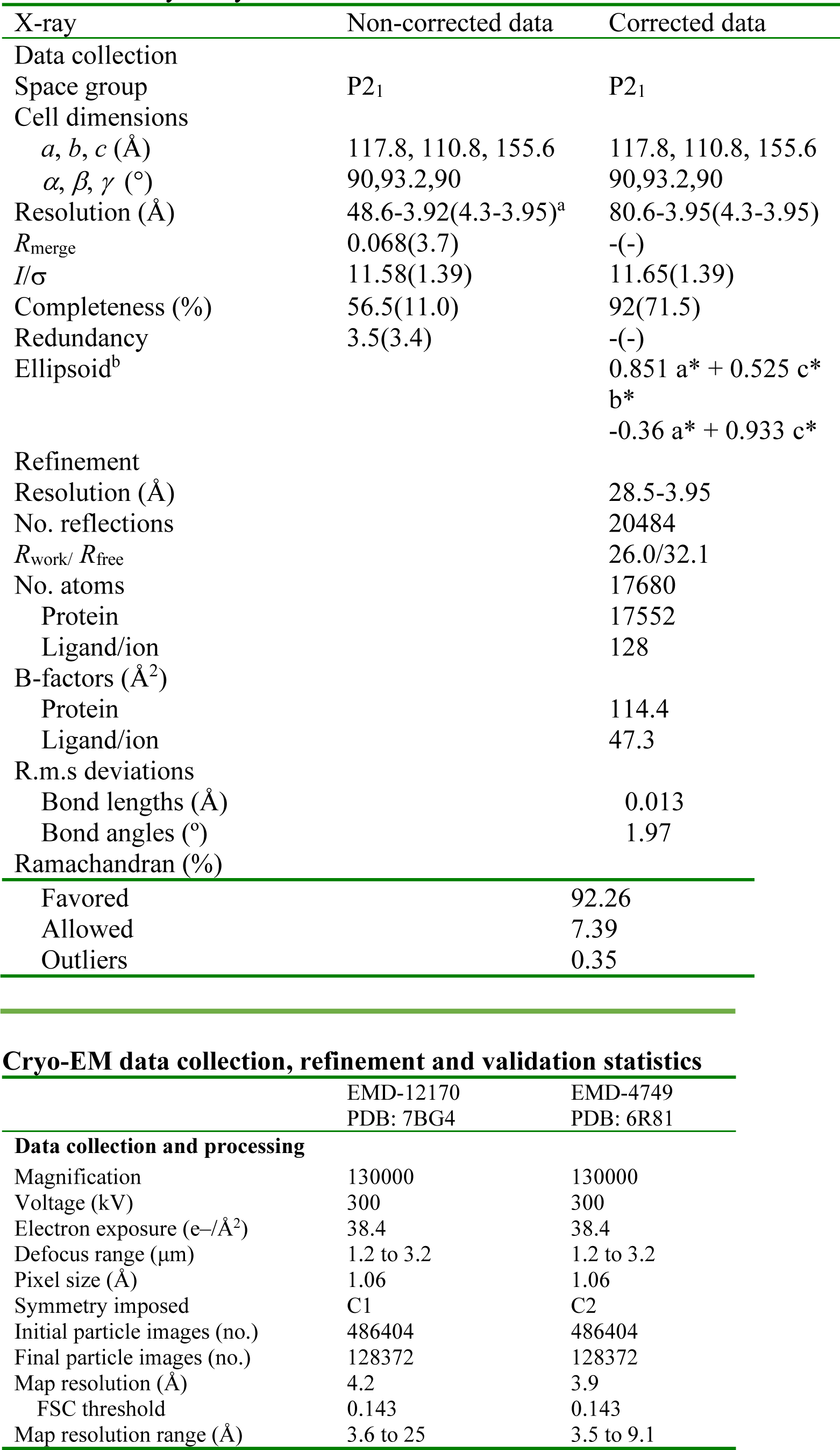
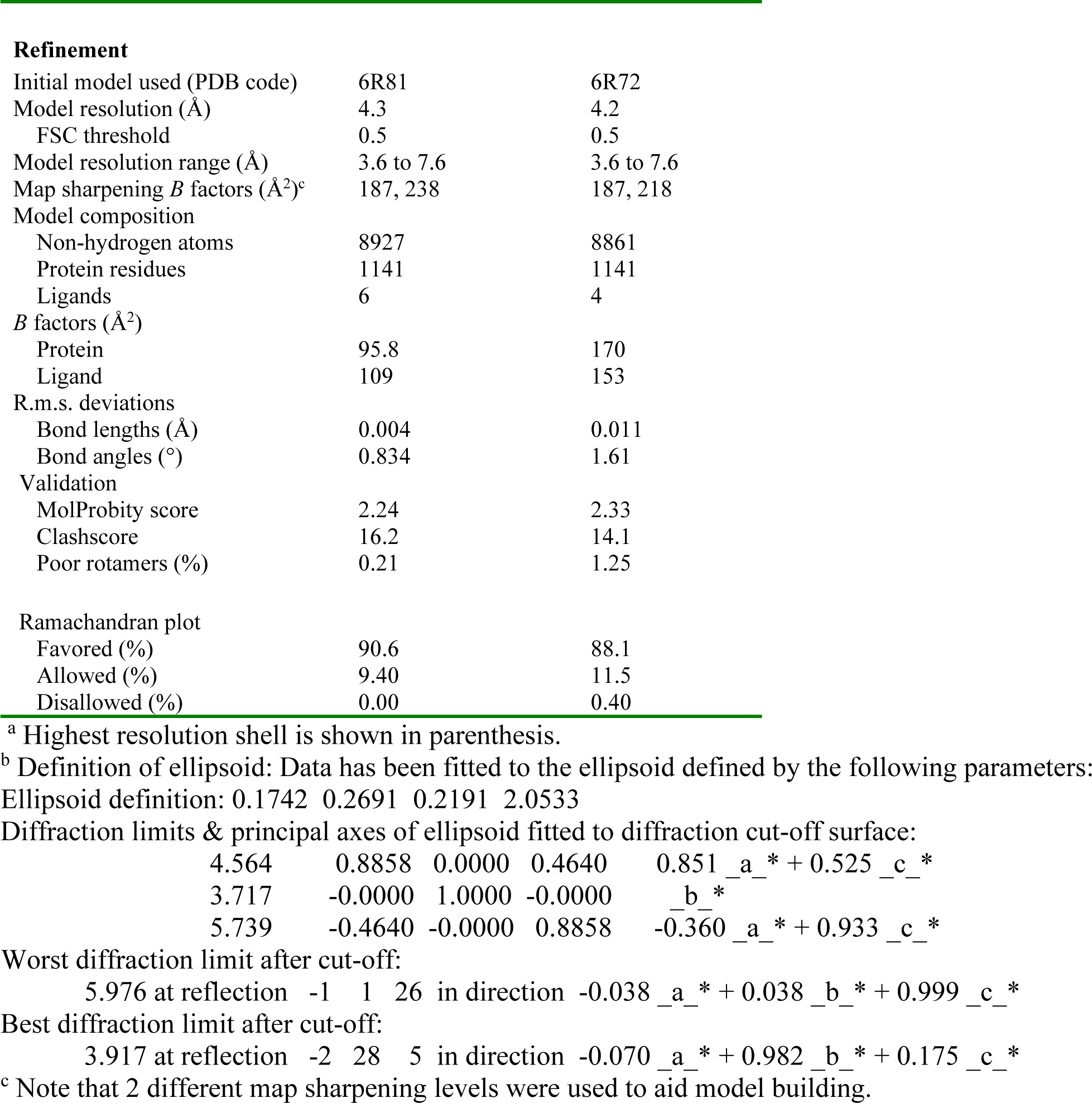
X-ray & cryo-EM data collection and refinement statistics

| X-ray | Non-corrected data | Corrected data |
| --- | --- | --- |
| Data collection |  |  |
| Space group | P2<sub>1</sub> | P2<sub>1</sub> |
| Cell dimensions |  |  |
| *a*, *b*, *c* (Å) | 117.8, 110.8, 155.6 | 117.8, 110.8, 155.6 |
| $\alpha$, $\beta$, $\gamma$ (°) | 90,93.2,90 | 90,93.2,90 |
| Resolution (Å) | 48.6-3.92(4.3-3.95)<sup>a</sup> | 80.6-3.95(4.3-3.95) |
| *R*<sub>merge</sub> | 0.068(3.7) | -(-) |
| *I*/$\sigma$ | 11.58(1.39) | 11.65(1.39) |
| Completeness (%) | 56.5(11.0) | 92(71.5) |
| Redundancy | 3.5(3.4) | -(-) |
| Ellipsoid<sup>b</sup> |  | 0.851 *a* * + 0.525 *c* *<br>*b* *<br>-0.36 *a* * + 0.933 *c* * |
| Refinement |  |  |
| Resolution (Å) |  | 28.5-3.95 |
| No. reflections |  | 20484 |
| *R*<sub>work</sub>/*R*<sub>free</sub> |  | 26.0/32.1 |
| No. atoms |  | 17680 |
| Protein |  | 17552 |
| Ligand/ion |  | 128 |
| B-factors (Å<sup>2</sup>) |  |  |
| Protein |  | 114.4 |
| Ligand/ion |  | 47.3 |
| R.m.s deviations |  |  |
| Bond lengths (Å) |  | 0.013 |
| Bond angles (°) |  | 1.97 |
| Ramachandran (%) |  |  |
| Favored |  | 92.26 |
| Allowed |  | 7.39 |
| Outliers |  | 0.35 |

**Cryo-EM data collection, refinement and validation statistics**
|  | EMD-12170<br>PDB: 7BG4 | EMD-4749<br>PDB: 6R81 |
| --- | --- | --- |
| **Data collection and processing** |  |  |
| Magnification | 130000 | 130000 |
| Voltage (kV) | 300 | 300 |
| Electron exposure (e-/Å<sup>2</sup>) | 38.4 | 38.4 |
| Defocus range (μm) | 1.2 to 3.2 | 1.2 to 3.2 |
| Pixel size (Å) | 1.06 | 1.06 |
| Symmetry imposed | C1 | C2 |
| Initial particle images (no.) | 486404 | 486404 |
| Final particle images (no.) | 128372 | 128372 |
| Map resolution (Å) | 4.2 | 3.9 |
| FSC threshold | 0.143 | 0.143 |
| Map resolution range (Å) | 3.6 to 25 | 3.5 to 9.1 |
| **Refinement** |  |  |
| Initial model used (PDB code) | 6R81 | 6R72 |
| Model resolution (Å) | 4.3 | 4.2 |
| FSC threshold | 0.5 | 0.5 |
| Model resolution range (Å) | 3.6 to 7.6 | 3.6 to 7.6 |
| Map sharpening *B* factors (Å<sup>2</sup>)<sup>c</sup> | 187, 238 | 187, 218 |
| Model composition |  |  |
| Non-hydrogen atoms | 8927 | 8861 |
| Protein residues | 1141 | 1141 |
| Ligands | 6 | 4 |
| *B* factors (Å<sup>2</sup>) |  |  |
| Protein | 95.8 | 170 |
| Ligand | 109 | 153 |
| R.m.s. deviations |  |  |
| Bond lengths (Å) | 0.004 | 0.011 |
| Bond angles (°) | 0.834 | 1.61 |
| Validation |  |  |
| MolProbity score | 2.24 | 2.33 |
| Clashscore | 16.2 | 14.1 |
| Poor rotamers (%) | 0.21 | 1.25 |
| Ramachandran plot |  |  |
| Favored (%) | 90.6 | 88.1 |
| Allowed (%) | 9.40 | 11.5 |
| Disallowed (%) | 0.00 | 0.40 |

|  |  |  |  |  |
| --- | --- | --- | --- | --- |
| 4.564 | 0.8858 | 0.0000 | 0.4640 | $0.851 \text{ } _a \text{ } ^* + 0.525 \text{ } _c \text{ } ^*$ |
| 3.717 | -0.0000 | 1.0000 | -0.0000 | $\text{ } _b \text{ } ^*$ |
| 5.739 | -0.4640 | -0.0000 | 0.8858 | $-0.360 \text{ } _a \text{ } ^* + 0.933 \text{ } _c \text{ } ^*$ |

Worst diffraction limit after cut-off:
5.976 at reflection -1 1 26 in direction $-0.038 \text{ } _a \text{ } ^* + 0.038 \text{ } _b \text{ } ^* + 0.999 \text{ } _c \text{ } ^*$
Best diffraction limit after cut-off:
3.917 at reflection -2 28 5 in direction $-0.070 \text{ } _a \text{ } ^* + 0.982 \text{ } _b \text{ } ^* + 0.175 \text{ } _c \text{ } ^*$
<sup>c</sup> Note that 2 different map sharpening levels were used to aid model building.

**Data S1.**
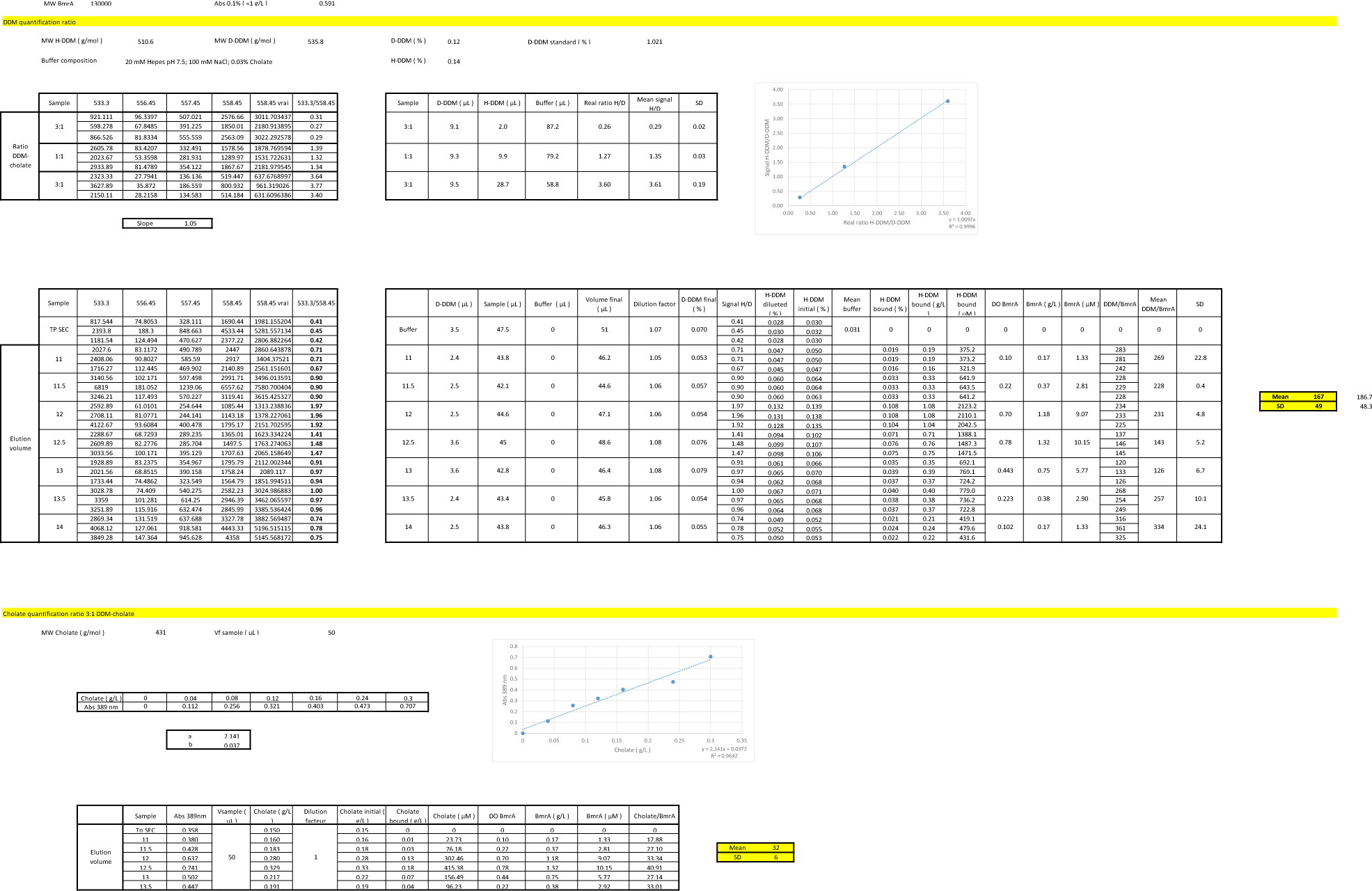
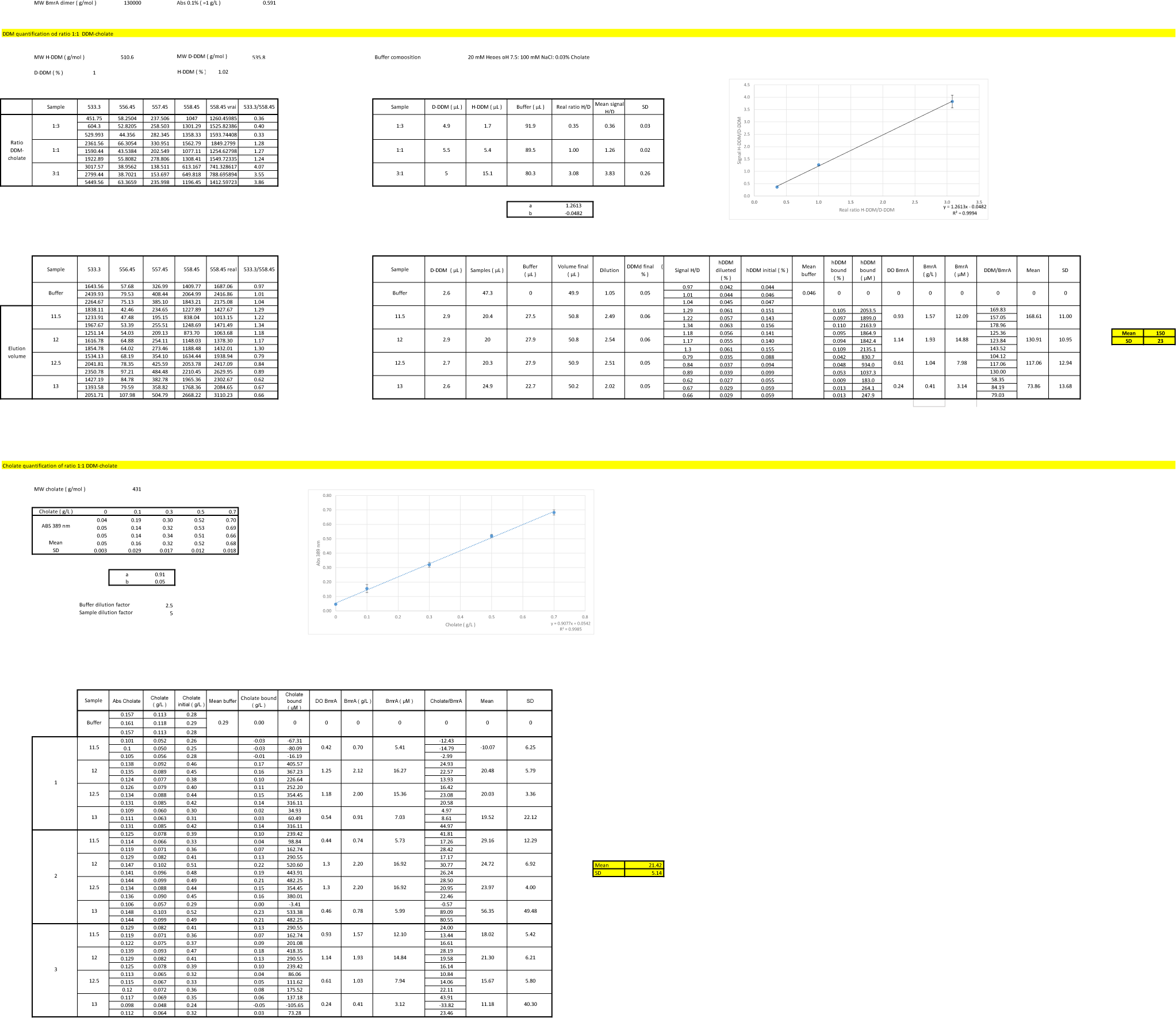
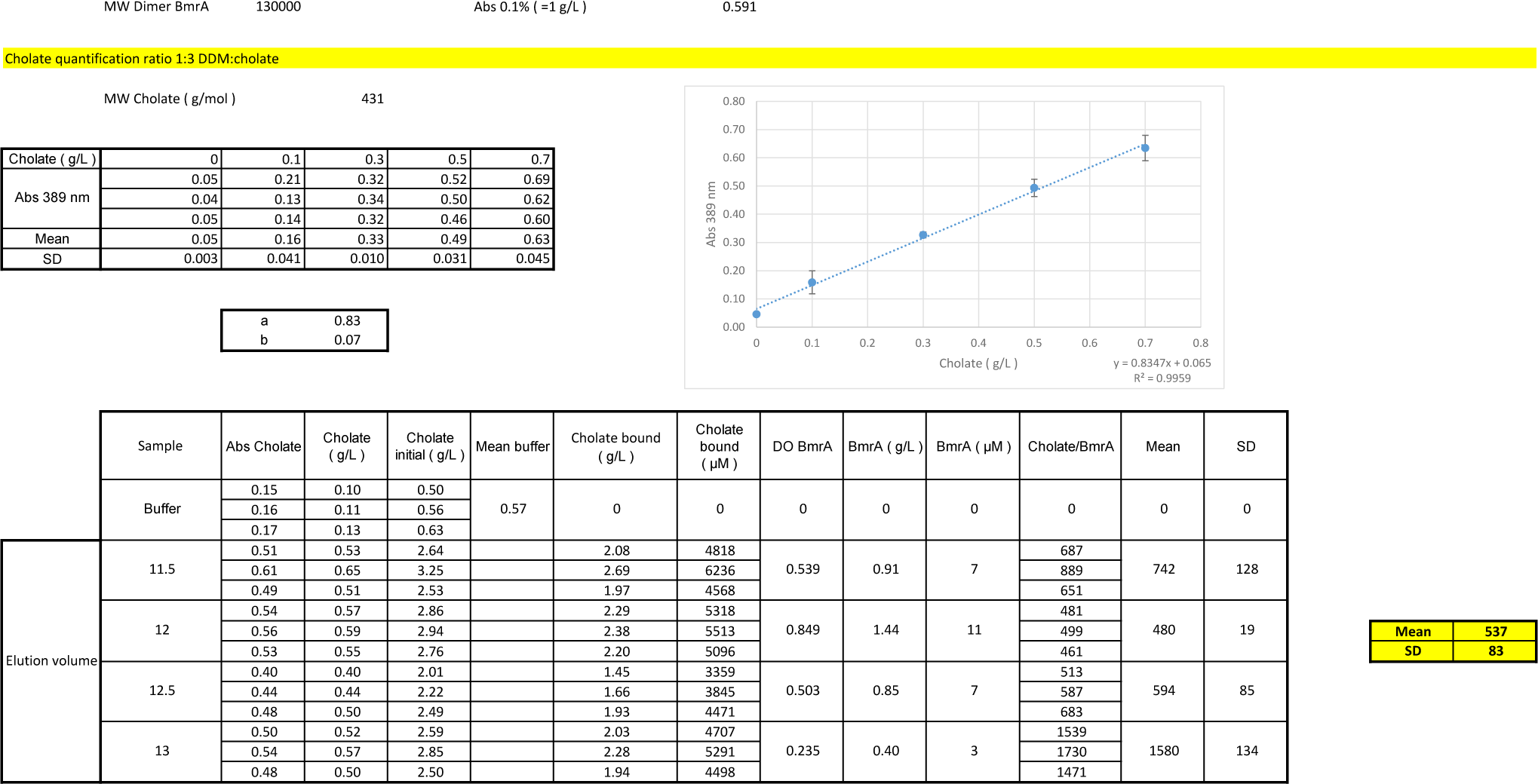
(separate file) Detergent quantitation.

**Data S2.**
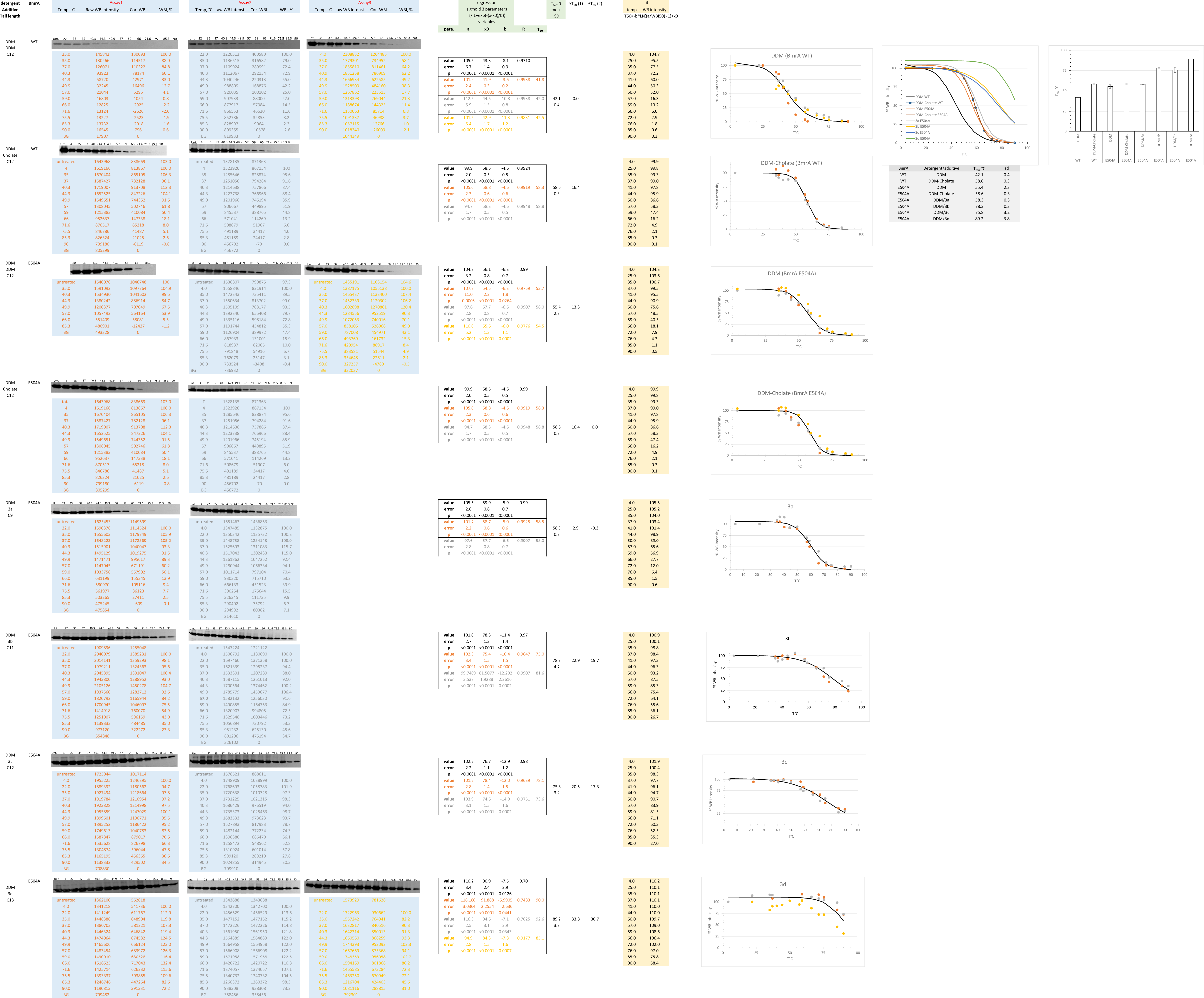
(separate file) Thermostability assays.

